# A novel splicing graph allows a direct comparison between exon-based and splice junction-based approaches to alternative splicing detection

**DOI:** 10.1101/2024.05.20.595048

**Authors:** Jelard Aquino, Daniel Witoslawski, Steve Park, Jessica Holder, Amei Amei, Mira V. Han

## Abstract

There are primarily two computational approaches to alternative splicing detection: splice junction-based and exon-based approaches. Despite their shared goal of addressing the same biological problem, these approaches have not been reconciled before. We devised a novel graph structure and algorithm aimed at mapping between the exonic parts and splicing events detected by the two different methods. Through simulations, we demonstrated disparities in sensitivity and specificity between splice junction-based and exon-based methods. When applied to empirical data, there were large discrepancies in the results, suggesting that the methods are complementary. With the discrepancies localized to individual events and exonic parts, we were able to gain insights into the strengths and weaknesses inherent in each approach. Finally, we integrated the results to generate a comprehensive list of both common and unique alternative splicing events detected by both methodologies.

**Availability:** https://github.com/HanLabUNLV/GrASE

**Supplementary information:** Supplementary data are available online.

## 1. Introduction

Alternative splicing (AS) enhances the transcriptomic and proteomic diversity in eukaryotic cells by generating different isoforms through the inclusion or exclusion of specific exons. The AS process is involved in essential biological processes, such as development[1], differentiation[2], and disease[3–5]. Computational tools analyzing alternative splicing in RNA-seq data broadly fall into two categories: splice junction-based and exon-based approaches. The former identifies AS events by analyzing junction reads spanning splice sites, while the latter analyzes reads mapping to exon fragments. Popular tools like rMATS[6], SUPPA2[7], and MAJIQ[8] use the splice junction-based approach while others like DEXSeq[9] and JunctionSeq[10] use the exon fragment-based approach. While the splice junction-based approach has a more direct correspondence to specific alternative splicing events, it relies on fewer reads that map to fewer bases at the junctions, potentially leading to lower power in case of low coverage or low expression events. The exon-based approach considers all reads overlapping the exon of interest, including those not spanning a splice junction, so it can potentially lead to higher power relying on more coverage, but because it exhaustively tests every exonic part in the gene, unaware of the splicing structure, it leads to larger numbers of tests and suffers more from the multiple testing problem [11]. Also, the exonic part that is the unit of the exon-based approach can sometimes be involved in multiple alternative splicing events that are nested among each other, or even alternative transcription start (ATS) or alternative transcription termination (ATT) events, so there may not be a direct one-to-one relationship between the splicing event vs. the exonic part usage.

Despite these complementary strengths and weaknesses, no studies have directly compared these approaches at the event and exon level. There have been several studies that systematically and comprehensively compared multiple AS software. But, because it is not straight-forward to map the ‘events’ of event-based tools to the ‘exonic parts’ of exon-based tools, usually the problem is circumvented by avoiding the direct comparison at the event/exon level. The studies either ignore exon-based methods and only compare the event-based methods [12–14] or they aggregate the results to the gene level. The aggregation from event/exon level results to the gene level results are done by either limiting the simulation or empirical data to genes that have exactly two transcripts and only a single splicing event [15], or by summarizing the *p-values* of all the events identified for each gene into a single *q-value* per gene [11,16]. To bridge this gap between the splice junction-based and exon fragment-based approach, we present a novel splicing graph, GrASE (Graph of Alternative Splice junctions and Exonic parts), that produces a mapping between exonic parts and the corresponding splicing events. The capability to map between the two approaches allows two important analyses that were not available before. First, for the simple AS events where there is direct mapping between the two, we can compare and evaluate the performance of the two approaches at the event level. Second, we can compare the extent of consistency and complementarity of the two approaches against empirical data. Using a comprehensive simulated data, for the first time, we evaluated the performance of two representative software for each approach, rMATS and DEXSeq, for their specificity and sensitivity at the splicing event/exon level. Also, using cell-line and blood samples, we show how we can integrate the results from the two software and find the events that are common and unique.

## 2. Methods

### 2.1 Overview of GrASE

We introduce GrASE, a novel splicing graph. GrASE, a directed acyclic graph, includes splice junctions as nodes and three different types of edges: exonic parts, exons and introns. Exonic parts represent the exon counting bins, that are defined by flattening the exon boundaries derived from gene annotations using *dexseq_prepare_annotation*.*py* of DEXSeq [9]. The main difference between traditional splice graphs and GrASE are these exonic parts edges (*E*_*ex*_*part*_) that are arranged sequentially along genomic positions without overlap. GrASE resembles the contiguous splice graph introduced in Whippet [17], but GrASE represents junctions as nodes and exonic parts as edges, integrating the exonic parts edges with edges for regular exons and introns. Below we describe the structure of GrASE with an example graph G. The graph G comprises five nodes representing the five splice junctions, plus artificial start (*R*) and end (*L*) nodes. The edges consist of three disjoint set of edges, the intron edges (*E*_*intron*_), the exon edges (*E*_*exon*_) and the exonic part edges (*E*_*ex*_*part*_) that connect the junctions based on the gene annotation.

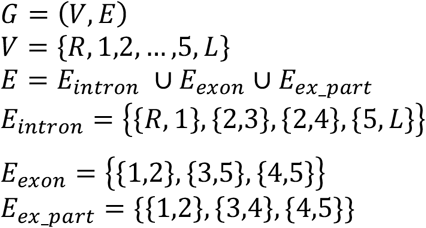

We also use the notation *intron*_*R*,1_ instead of {*R*, 1} ∈ *E*_*intron*_ for simplicity.

On this splicing graph, one can represent a transcript as i) a path along the exon and intron edges similar to traditional splicing graphs, or ii) a path along the exonic parts and intron edges. For example, in Figure 1, the same transcript ENST00000446393 can be represented as both

**Figure 1.**
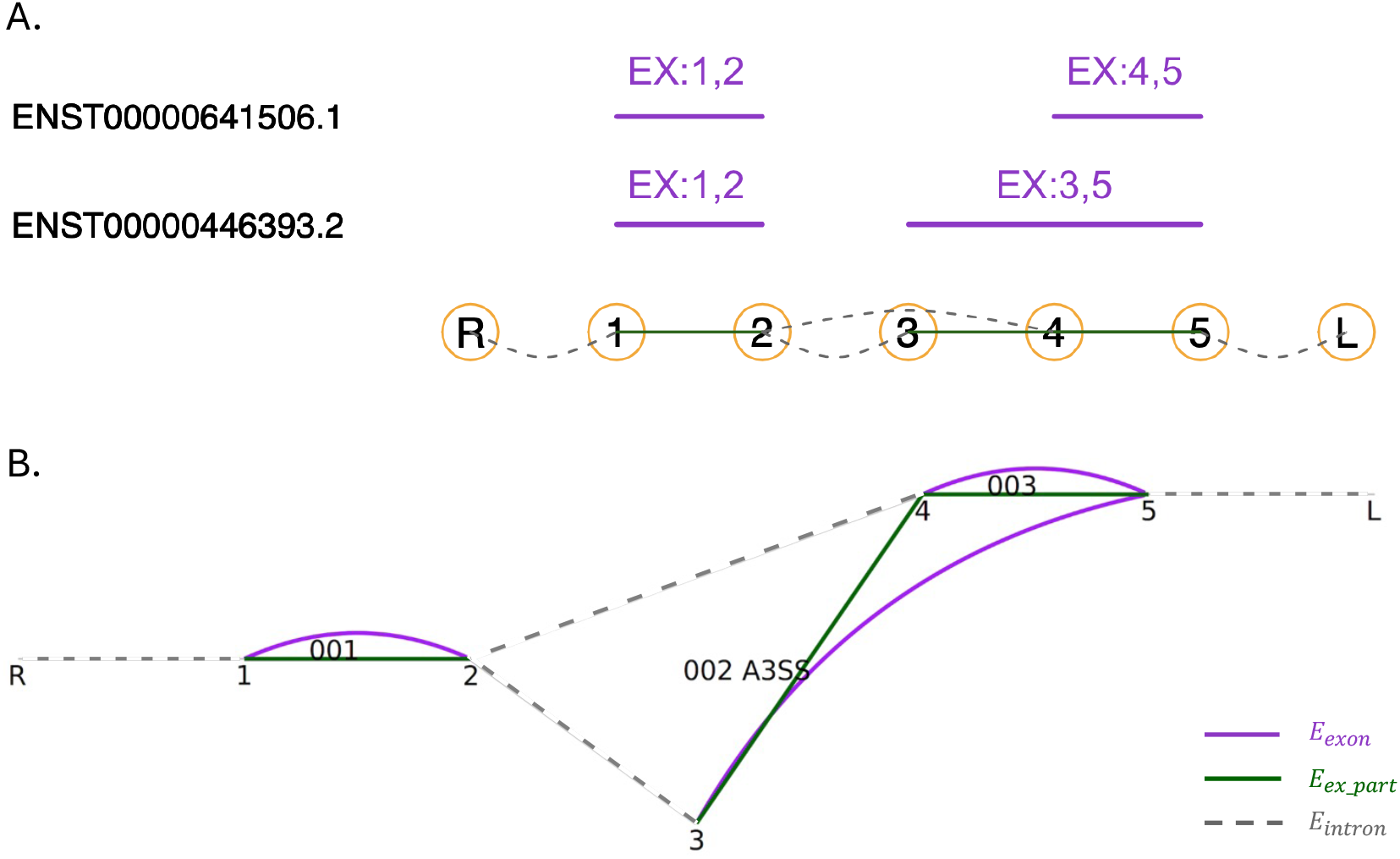
A) The transcript structure of the two transcript variants of gene OR9H1P (ENSG00000228336). B) GrASE graph representation of the gene. The nodes are the splice junctions, the intron edges (*E*_*intron*_) are represented by the dotted gray lines, the exon edges (*E*_*exon*_) are represented by the solid purple lines and the exonic part edges (*E*_*ex_part*_) are represented by the solid green lines. The exonic part edges are also labeled by the DEXSeq exonic part ID. We can see the splicing event identified as a bubble from node 2 to node 5, and the differential exonic part, *ex_part*_3,4_ (DEXSeq ID 002) that maps to the splicing event marked with the label A3SS.

i. *S*_ENST00000446393_ = {*intron*_*R*,1_, *exon*_1,2_, *intron*_2,3_, *exon*_3,5_, *intron*_5,*L*_} along the intron/exon edges,
ii. *S*’_ENST00000446393_ = {*intron*_*R*,1_, *ex_part*_1,2_, *intron*_2,3_, *ex_part*_3,4_, *ex_part*_4,5_, *intron*_5,*L*_} along the intron/exonic part edges.

Likewise for the alternative transcript ENST00000641506, we have

i. *S*_ENST00000641506_ = {*intron* _*R*,1_, *exon*_1,2_, *intron*_2,3_, *exon*_4,5_, *intron*_5,*L*_} along the intron/exon edges,
ii. *S*^′^_ENST00000641506_ = {*intron* _*R*,1_, *ex_part*_1,2_, *intron*_2,3_, *ex_part*_4,5_, *intron*_5,*L*_} along the intron/exonic part edges.

The unique property of exonic part edges lies in their one-dimensional genomic arrangement, defined by flattened exon boundaries, ensuring non-overlapping intervals. Consequently, for any exon defined by start and end nodes, there exists a single unique path in set *E*_*ex*_*part*_. This guarantees a one-to-one mapping between exon edges (*exon*_*i,j*_) and corresponding paths through exonic part edges. Examples in Figure 1 are the mapping from the exon edges (*exon*_3,5_) to exonic part edges (*ex_part*_3,4_, *ex_part*_4,5_), and the mapping from the exon edges (*exon*_4,5_) to exonic part edges (*ex_part*_4,5_).

A splicing event manifests as a bubble in the splicing graph, formed by splits where a node defining an exonic part edge has multiple outgoing/incoming degree of exon or intron edges [18]. To detect alternative splicing, we focus on nodes connected with multiple edges, as these represent potential junctions for AS. In Figure 1B, node 2 splits into two outgoing intron edges and node 5 splits into two incoming exon edges. Those alternative paths create bubbles in the splicing graph and define an alternative splicing event as a set of subpaths {*s*_1_, *s*_2_} that start at the same node (node 2) and end at the same node (node 5).

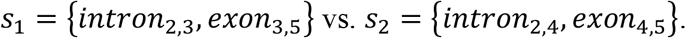

To find the mapping between the exonic part and the splicing event, we leverage the previously described mapping between the exon paths and exonic part paths. For a splicing event that is defined as two alternative paths along the intron/exon edges, we identify the corresponding paths along the intron/exonic part edges.

In Figure 1B, the corresponding alternative subpaths would be 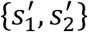,

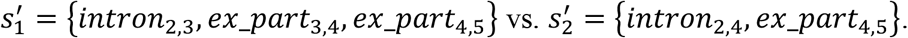

We translate these alternative paths along the intron/exonic parts 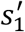 and 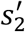 into sets. Exonic parts not common between the two paths represent the exonic parts used differently depending on the isoform usage. Thus, we find the symmetric difference between the two sets, and the exonic part edges that are remaining in the symmetric difference are the focal exonic parts that should be tested for differential exon usage for the splicing event 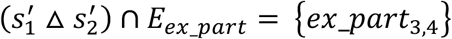.

We have now defined a mapping between the alternative splicing event {*s*_1_, *s*_2_} and the exonic parts 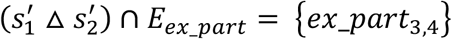.

### 2.2 Construction of GrASE and mapping of events to exonic parts

We constructed GrASE for all the gene models annotated in the human reference genome (Gencode v34). Using *dexseq_prepare_annotation*.*py* from DEXSeq [9], we identified all the exonic part boundaries. We used the SplicingGraphs R package [19] to construct the basic splicing graph consisting of introns and exons. We exported the basic graph as a graphml object and incorporated all the exonic part edges defined by DEXSeq using the iGraph package in python.

While the concept of AS as bubbles in the graph seems straightforward in theory, detecting and classifying them into different AS types in practice can be challenging, particularly due to nested bubbles. For a long time, many software tools only addressed simple cases and ignored complex events. An alternative approach is to disregard the whole path defining the bubbles, and focus solely on the source and sink nodes separately. Then, you can view AS as two separate events: one defined by alternative outgoing edges for the source node, and the other defined by alternative incoming edges for the sink node. This approach was adopted by Vaquero-Garcia et al. [8] with their Local Splicing Variations.

Since the identification and classification of events are beyond the scope of our paper, we side-stepped on this issue, and relied on the splicing events that are empirically identified in a given RNAseq dataset by rMATS and DEXSeq for downstream analyses. The initial mapping was done starting from the four type of events (A3SS, A5SS, RI, and SE) found by rMATS in their *fromGTF*.*[AS]*.*txt* files, which were mapped to the exonic fragments defined by DEXSeq using the algorithm described in the previous section. Subsequently, we merged the mapping retrieved for all the rMATS events with the DEXSeq results.

### 2.3 Simulation of differential isoform expression

For our simulations, we selected three genes, each containing at least two splicing events positioned distantly within the gene structure, to highlight the functionality of splice-junction/exon level mapping. MPL contained an A3SS event at exon 3 and a SE event at exon 14; RBBP5 contained an A5SS event at exon 2 and a SE event at exon 15; TRERF1 contained an A3SS event at exon 6 and two A5SS events at exon 15 and exon 23 (Supplementary Figure S1). For TRERF1, we focused on the A3SS and the second A5SS event annotated in Supplementary Figure S1. We used Polyester [20] to simulate raw RNA-seq reads for five samples each across two conditions. Polyester models isoform-level differential expression across biological replicates. For each event, we manipulated a randomly chosen single transcript encompassing the alternative splicing event to have the fold change and read depth parameters described below. In total, we generated 500,000 (2 conditions x 5 samples x 100 fold change x 500 read depths) sets of simulated RNA-seq data for each event, covering all combinations of read depth and fold change parameters.

To determine simulation parameters, we analyzed five replicates for B naive cells and five replicates for CD8 naive cells from Bonnal et al. (Accession ID: PRJEB5468) [26]. Based on the estimated distribution of log2 Fold Change between the two cell types (Supplementary Figure S2), we designed the fold change parameters seen in Supplementary Table 1. Based on the estimated distribution of read counts mapped to transcripts for each sample (Supplementary Figure S3), we designed a similar read depth per transcript distribution for the simulation (Supplementary Table 2). Here the *read depth* is used as a multiplier to determine the average number of reads per transcript, using the formula *read depth ∗ lenght of transcript* / *lenght of read* in our simulation. For the dispersion parameter we used two values representing large dispersion (smaller variance) and small dispersion (larger variance). For the large dispersion we used the default size parameter in Polyester [20] (1*/*3 of the transcript’s mean), and we used a value about 10-fold smaller (1*/*30 of the transcript’s mean) for the small dispersion.

### 2.4 Performance evaluation on simulated splicing events

The simulated reads were aligned using STAR against the particular gene region of the human reference genome (Gencode v34). Splice junction counts (JC) and splice junction exon counts (JCEC) were produced using rMATS (rmats.py), while exonic part counts were produced using DEXSeq. For each event and each sample, we combined the counts from each of the 50,000 simulated runs to create a synthetic “genome” with 50,000 “genes”, where each gene corresponds to a particular simulation with a specific fold change and read depth. This aggregation resulted in a single count matrix for each event, containing 50,000 rows and 10 columns for the 10 simulated samples. We applied rMATS’s statistical model on the JC and JCEC counts with the “—task stat” option. We ran DEXSeq on the combined exonic part counts.

For each simulated event, we utilized GrASE to identify the corresponding exonic parts mapping to the AS event. We compared the adjusted p-values provided by DEXSeq for the mapped exonic part to the FDR values provided by rMATS for the splicing event. The results based on the raw p-values before FDR correction are also reported for comparison. Simulations with fold change = 0 were considered real Negatives, while simulations with fold change *≠* 0 were considered real Positives for calculating sensitivity and specificity.

### 2.5 Comparison of rMATS vs. DEX-seq on empirical data

To compare results on empirical data, we analyzed AS between RNAseq of purified naive B cells and CD8+ naive T cells (Accession ID: PRJEB5468) [26]. The data was aligned against GRCh38 and Gencode v34 using STAR. We applied rMATS and DEXseq to quantify the counts and detect AS. Additionally, we compared RNA-seq on cell lines Raji and Jurkat for further comparison of homogenous samples.

## 3 Results

### 3.1 Direct comparison of the individual AS events detected by rMATS and DEXSeq on simulated data show higher sensitivity for rMATS and higher specificity for DEXSeq

Using GrASE, we were able to map the events and exonic parts in our simulation and compare the detection. Here, we present the results from the small dispersion (high variance) simulations, while results for large dispersion are available in supplementary figures (Supplementary Figures S4-S6). Sensitivity plots in Figures 2-4 illustrate a pattern where rMATS outperforms DEXSeq in detecting true positives, especially at lower read depths. The sensitivity correlates with read depth and fold change as expected. However, there is variation in the overall sensitivity across the same type of events in different genes. This variation is more pronounced for DEXseq, and is not associated with the length of the exonic part as hypothesized. Instead, it follows an unpredictable pattern dependent on which transcript was chosen for over/under expression, whether the exonic part was contained in the chose transcript, and the length of the chosen transcript. We describe this in more detail in section 3.2. This unpredictability affects the sensitivity of DEXSeq more severely, but has little effect on rMATS regardless of whether we are counting JC’s or JCEC’s.

**Figure 2.**
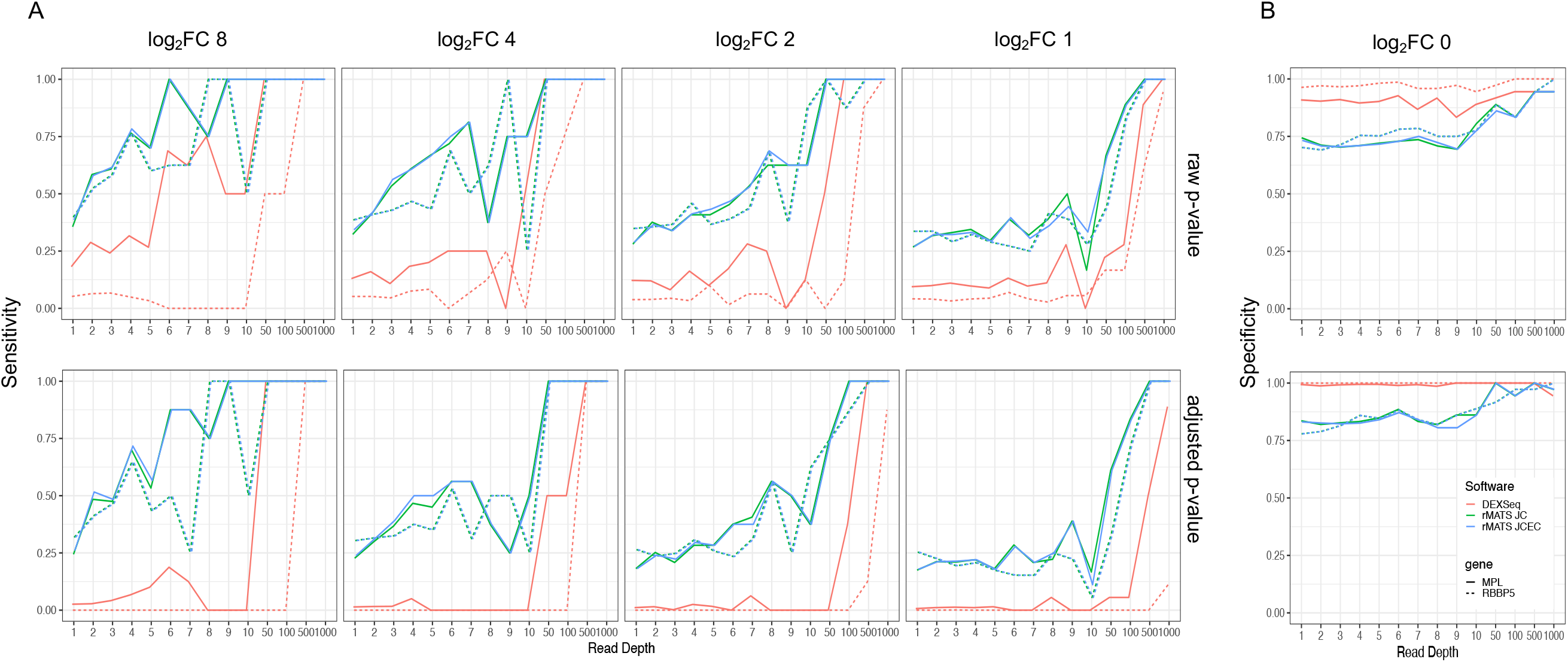
Sensitivity and specificity of skipped exon (SE) simulations at dispersion (σ) ≈ 0.033. **A)** Sensitivity results of the SE event in genes MPL (solid line) and RBBP5 (dotted line). Each panel represents the sensitivity (y-axis) vs the read depth (x-axis) for different log2 fold changes (8,4,2,1). The green line represents the performance based on rMATS’s junction counts (JC), the blue line rMATS’s junction and exon counts (JCEC), and the red line DEXSeq. **B)** Specificity results (log2 FC 0) of the SE event in MPL and RBBP5. The top panels represent the metric based on the raw p-value of the test, while the bottom panel represents the adjusted p-value.

**Figure 3.**
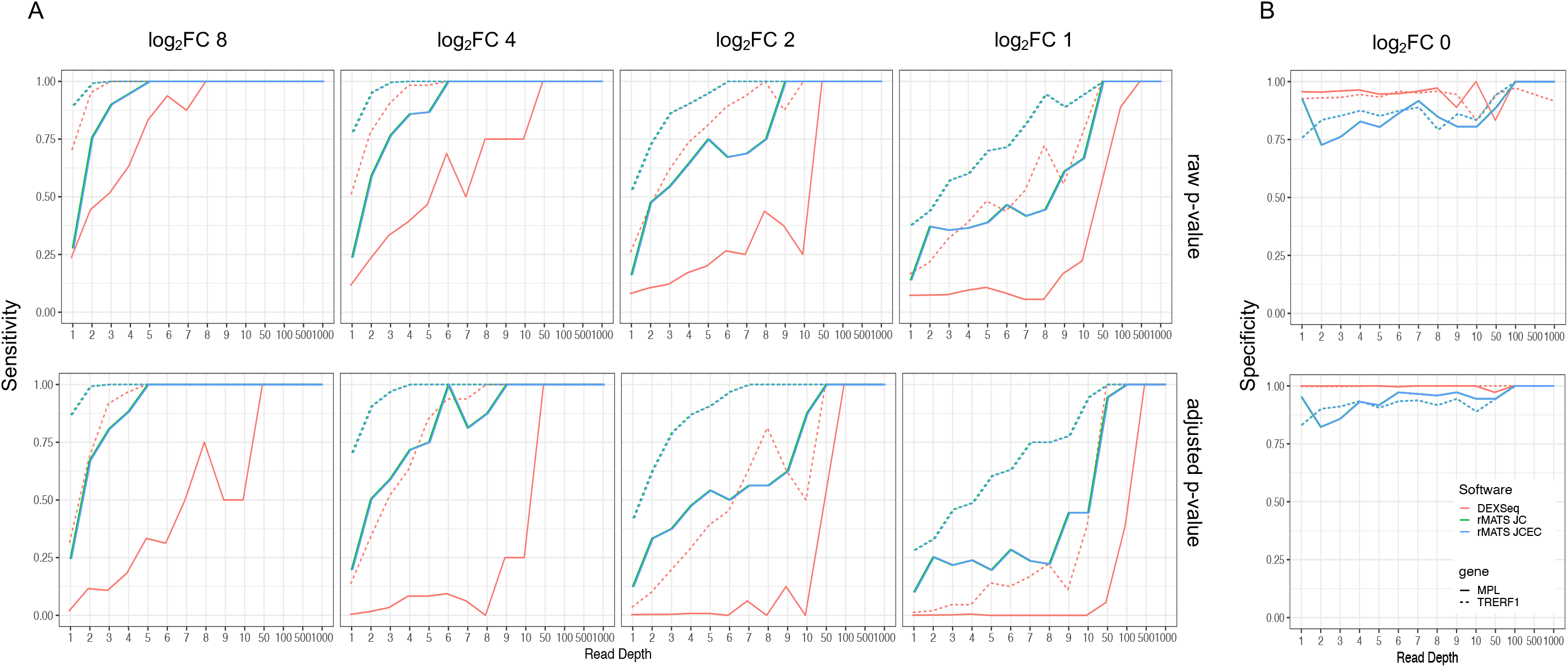
Sensitivity and specificity of alternative 3’ splice site (A3SS) simulations at dispersion (σ) ≈ 0.033. **A)** Sensitivity results of the A3SS event in genes MPL (solid line) and TRERF1 (dotted line). Each panel represents the sensitivity (y-axis) vs the read depth (x-axis) for different log2 fold changes (8,4,2,1). The green line represents the performance based on rMATS’s junction counts (JC), the blue line rMATS’s junction and exon counts (JCEC), and the red line DEXSeq. **B)** Specificity results (log2 FC 0) of the A3SS event in MPL and TRERF1. The top panels represent the metric based on the raw p-value of the test, while the bottom panel represents the adjusted p-value.

**Figure 4.**
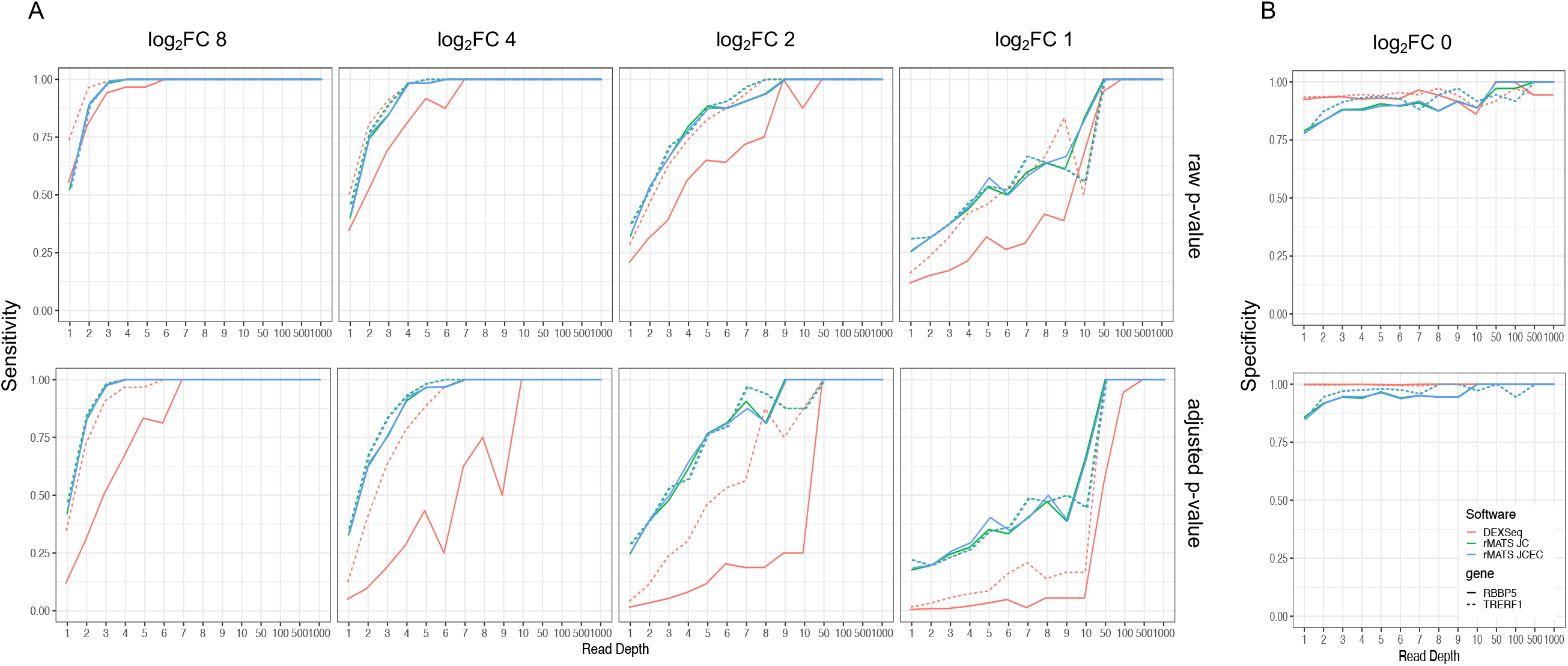
Sensitivity and specificity of alternative 5’ splice site (A5SS) simulations at dispersion (σ) ≈ 0.033. **A)** Sensitivity results of the A3SS event in genes RBBP5 (solid line) and TRERF1 (dotted line). Each panel represents the sensitivity (y-axis) vs the read depth (x-axis) for different log2 fold changes (8,4,2,1). The green line represents the performance based on rMATS’s junction counts (JC), the blue line rMATS’s junction and exon counts (JCEC), and the red line DEXSeq. **B)** Specificity results (log2 FC 0) of the A5SS event in RBBP5 and TRERF1. The top panels represent the metric based on the raw p-value of the test, while the bottom panel represents the adjusted p-value.

In contrast, the loss in sensitivity for DEXSeq was compensated by a higher specificity, especially in the small dispersion / high variance condition. We also observed that DEXSeq suffers more severely from multiple testing as the sensitivity drops drastically when the p-values are corrected. The observed pattern of performance was consistent across different isoform structures and different type of events.

This observed pattern is also reflected in accuracy metrics, with rMATS demonstrating higher accuracy at lower fold changes, and DEXSeq demonstrating higher accuracy at higher fold changes. As expected, DEXSeq shows relatively higher accuracy than rMATS in the smaller dispersions (Supplementary Figures S7–S9), and relatively lower accuracy than rMATS in the larger dispersions (Supplementary Figures S10– S12). Overall accuracy levels are high because simulations of a small number of positive cases with different fold changes and read depths are combined with simulations of a large number of negative cases with zero fold change to calculate the accuracy. This leads to negative cases dominating accuracy measures, and specificity is weighted more importantly than sensitivity. But we believe this better reflects the imbalanced empirical positive rate and is a more realistic scenario.

### 3.2 DEXSeq misidentifies significant exons in certain conditions

In our simulation, we encountered a specific case in the Skipped Exon of RBBP5 (Figure 2), where even at large fold changes of up to 8x and high read depths of 10 reads per base, DEXSeq failed to detect the event. Curious about this result, we investigated it further, and discovered that such occurrences arise when the combination of exon lengths and transcript length in the gene structure results in a consistent ratio between the “*this*” and “*others*” components, even after significant fold changes. In the case of RBBP5, the manipulated transcript includes the exonic part, and exhibited a large difference in overall length due to an alternative TTS. Due to a particular length relationship, the ratio between “*this*” and “*other*” ended up being similar despite a large fold change.

Since this case was encountered serendipitously, we sought to identify similar instances. Utilizing a condition where the ratio between “this” and “other” remains equal after a large fold change, we developed a formula (Supplementary Figure S13) that when minimized, allowed us to identify potential cases where even a large fold change wouldn’t lead to a significant interaction term. We selected 25 cases of gene structures with the smallest values for our condition formula based on the Gencode annotations, and simulated a favorable scenario of large fold change (8x) and high read depth (100) for these genes. Among the 25 cases, we found that DEXSeq could not detect the skipped exon in 8 instances (Supplementary Table 3). However, in all these cases, DEXSeq did identify significant exon usage difference in the adjacent exons neighboring the skipped exon. This limitation has been discussed in the original publication, wherein they noted that the test’s output with regard to which of the counting bins are affected can be unreliable if the isoform regulation affects a large fraction of the exons [21]. By directly mapping the splicing event to the relevant exon between rMATS and DEXSeq, we could detect this misidentification of the exonic part and confirm its impact even in cases with simple gene structure featuring few splicing events.

### 3.3 GrASE allows mapping between rMATS events and DEXSeq exonic parts

After examining simulated data, we compared the detection in empirical data. We conducted differential splicing analysis using rMATS and differential exon usage analysis using DEXSeq on RNA-seq of highly purified naive B and naive CD8^+^ T cells from Bonnal et al. [26]. For all alternative splicing (AS) events detected by rMATS, we successfully identified the corresponding exonic parts that differed between the alternative splicing events. Figure 5 illustrates three genes as examples, each exhibiting different kinds of splicing events (SE, A3SS, and A5SS respectively). Based on the Gencode v34 annotation, there were a total of 585,104 exonic parts. Comparing the naive B and naive T cells, rMATS detected 135,880 AS events, which mapped to 138,997 differential exonic parts using GrASE. Meanwhile, DEXSeq tested 308,737 exonic parts out of the 585,104 after independent filtering. Overall, there was a discrepancy in the events and exonic parts tested between the two software, with only 83,502 events (equivalent to 84,208 exonic parts) tested by both software. There was a considerable number of exonic parts that are tested by DEXSeq but were ignored by rMATS. Because DEXSeq tests all exonic parts that pass the independent filtering, the discrepancy in the total number of tests may arise from exonic parts that are not involved in any isoform switch, but also from exonic parts that are involved in isoform switch that rMATS does not consider. The latter includes 1) Alternative Transcript Start Site (TSS) or Alternative Transcription Termination Site (TTS) [22] that rMATS disregards, and 2) complex events that are missed by rMATS [8]. The gene with the highest number of events detected was the long non-coding RNA Linc-GALH, which had a total of 146 different events, encompassing 178 different exonic parts in various combinations. Six genes had more than 100 events detected and mapped to exonic parts, including *TEX41, Linc-GALH, PVT1, SPG7, SNHG14*, and *DDX3X*, four of which are long noncoding RNAs.

**Figure 5.**
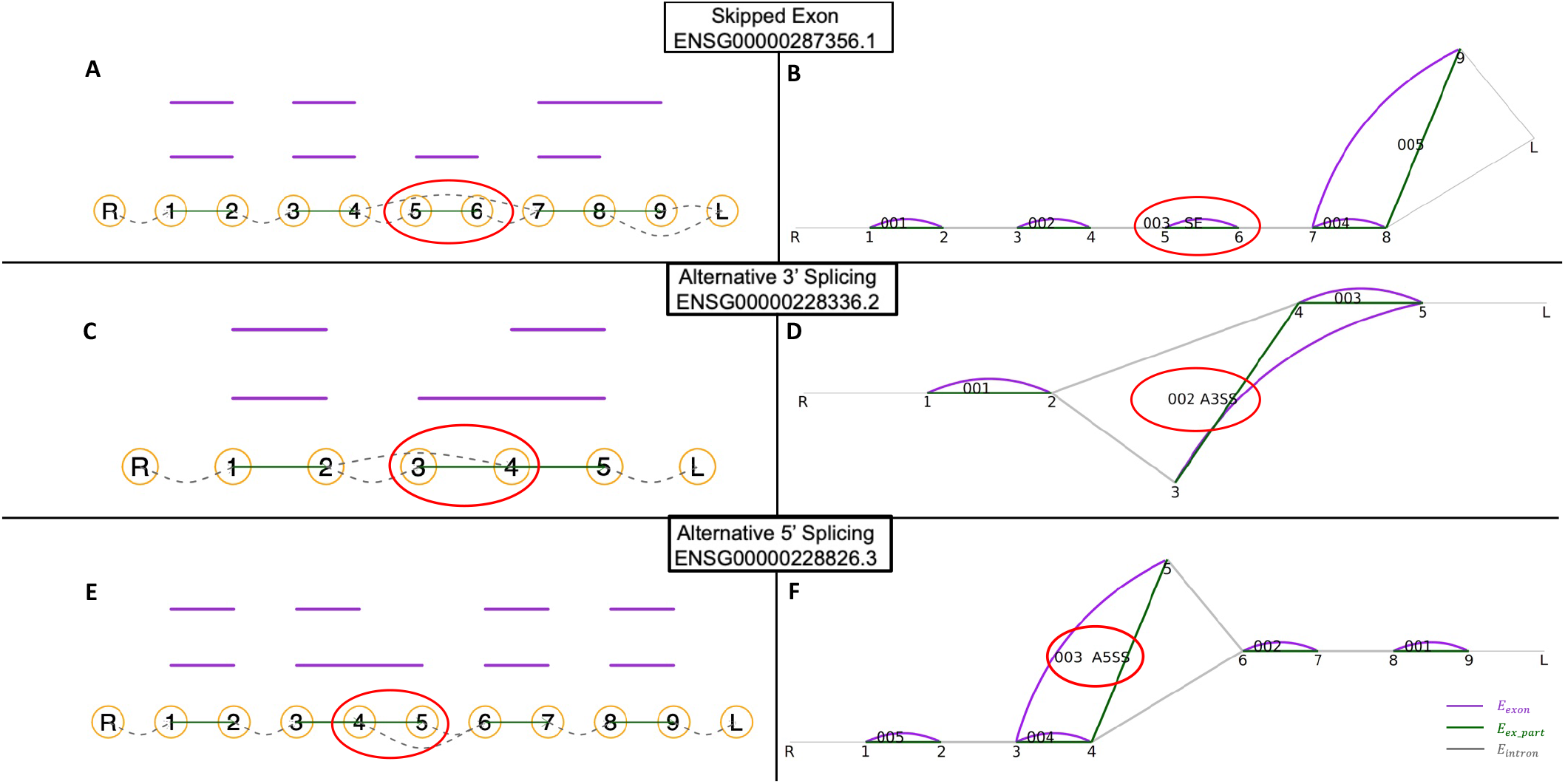
Mapping between the AS event and the focal exonic part. Panels on the left describe the AS event and the transcripts involved, while the right panels visualize GrASE and highlight the focal exonic part that is differentially included between the two alternative splice forms. A,B) ENSG00000287356 with a skipped exon (SE) event. C,D) ENSG00000228336 (*OR9H1P*) with an A3SS event. E,F) ENSG00000228826 with an A5SS event.

### 3.4 Direct comparison of the individual AS events detected by rMATS and DEXSeq on empirical data show large discrepancy

We identified 3,504 differential splicing events (FDR ≤ 0.05) through rMATS. For differential exon usage, we identified 3,647 exonic parts that were differentially used (adjusted p-value ≤ 0.05) based on DEXSeq. After running each software separately and comparing the results, we mapped the splicing events from rMATS to their corresponding the exonic parts from DEXSeq using GrASE, to examine the concordance between the two software. We found that the two software are largely complementary, with only a minor proportion of significant calls from each software showing common detection (Figure 6). Reporting on the intersection and union of events and exonic parts turned out to be slightly complicated due to the 1:*n* or *n*:1 mapping that exists between events and exonic parts, when a single event encompasses several exonic parts or when a single exonic part is involved in several different splicing events. So, here we report it in two different ways, first through the number of events, and second, through the number of exonic parts. To find the intersection, for cases of one-to-many mapping, if at least one of the many mapped events/exonic parts was called significant we considered it to be called significant. Based on the number of events, there were a total of 135,880 events that were identified and tested by rMATS and we found corresponding exonic parts to all of those cases. Of those, 268 were called significant with both rMATS and DEXSeq, 3,236 were only called with rMATS, and 1,205 were only called with DEXSeq. Based on the number of exonic parts, there were a total of 308,737 exonic parts tested with DEXSeq after independent filtering, and we mapped 138,997 of them to the corresponding events detected with rMATS. Of those tested with both rMATS and DEXSeq, 289 were called significant with both rMATS and DEXSeq, 7,143 were only called with rMATS, and 528 were only called with DEXSeq. Additionally, there were 100 exonic parts that were never tested by rMATS but called significant with DEXSeq.

**Figure 6.**
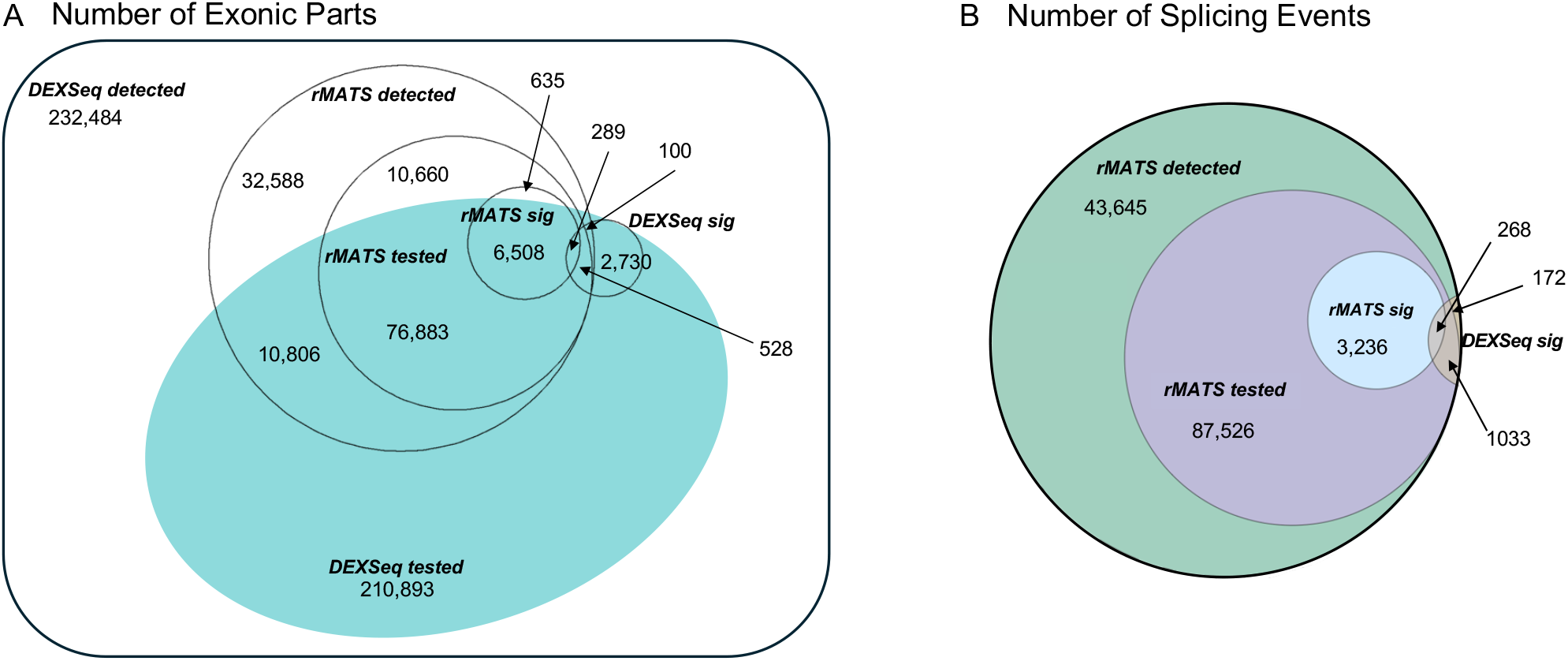
Venn diagrams for exonic parts and splicing events that are detected, tested and called positive. A) Overlapping in terms of DEXSeq exonic parts. B) Overlapping in terms of rMATS splicing events. rMATS detected = all of the splicing events annotated by rMATS; rMATS tested = all of the splicing events rMATS statistically tested; rMATS sig = all the significant splicing events in rMATS with FDR ≤ 0.05; DEXSeq detected = all the exonic parts annotated by DEXSeq; DEXSeq tested = all exons DEXSeq statistically tested that were given an adjusted p-value; DEXSeq sig = all the significant exons DEXSeq detected with adjusted p-value ≤ 0.05.

### 3.5 Reducing the number of tests DEXSeq runs to exons with potential AS events did not improve the performance for DEXSeq

As seen in our simulated results, DEXSeq exhibited lower sensitivity due to the large number of multiple tests it conducts for all the exons. We hypothesized that by reducing the number of tests that DEXSeq performs, we can boost the sensitivity and identify more exons involved in alternative splicing as differentially used. This idea was originally explored in Soneson et al. where they filtered out lowly expressed transcript isoforms to reduce the number of tests and demonstrated improved performance [11]. But, in practice, the true expression level of an isoform is unknown. Here, we adopted a different approach where instead of filtering the isoforms which we cannot directly count, we filtered the exonic parts. Leveraging the mapping between the exonic parts and rMATS, we can identify the exonic parts involved in the candidate splicing events that are identified by rMATS, and only test those relevant exonic parts. We tested this hypothesis by filtering the results of the DEXSeq tests to only retain the exonic parts involved in an AS event that is tested in rMATS. Subsequently, we applied DEXSeq’s multiple testing correction function, including the independent filtering to the retained list of relevant exonic parts. Although the total number of exons to be tested was reduced to about one fifth of the original list, it did not have the effect that we hypothesized. With the reduced list of relevant exonic parts, the independent filtering step of DEXSeq removed more exons from testing, but it also reduced the number of positive calls. In the end, the proportion of positives that DEXSeq detected among the relevant exonic parts were slightly lower than before filtering. (Figure 7).

**Figure 7.**
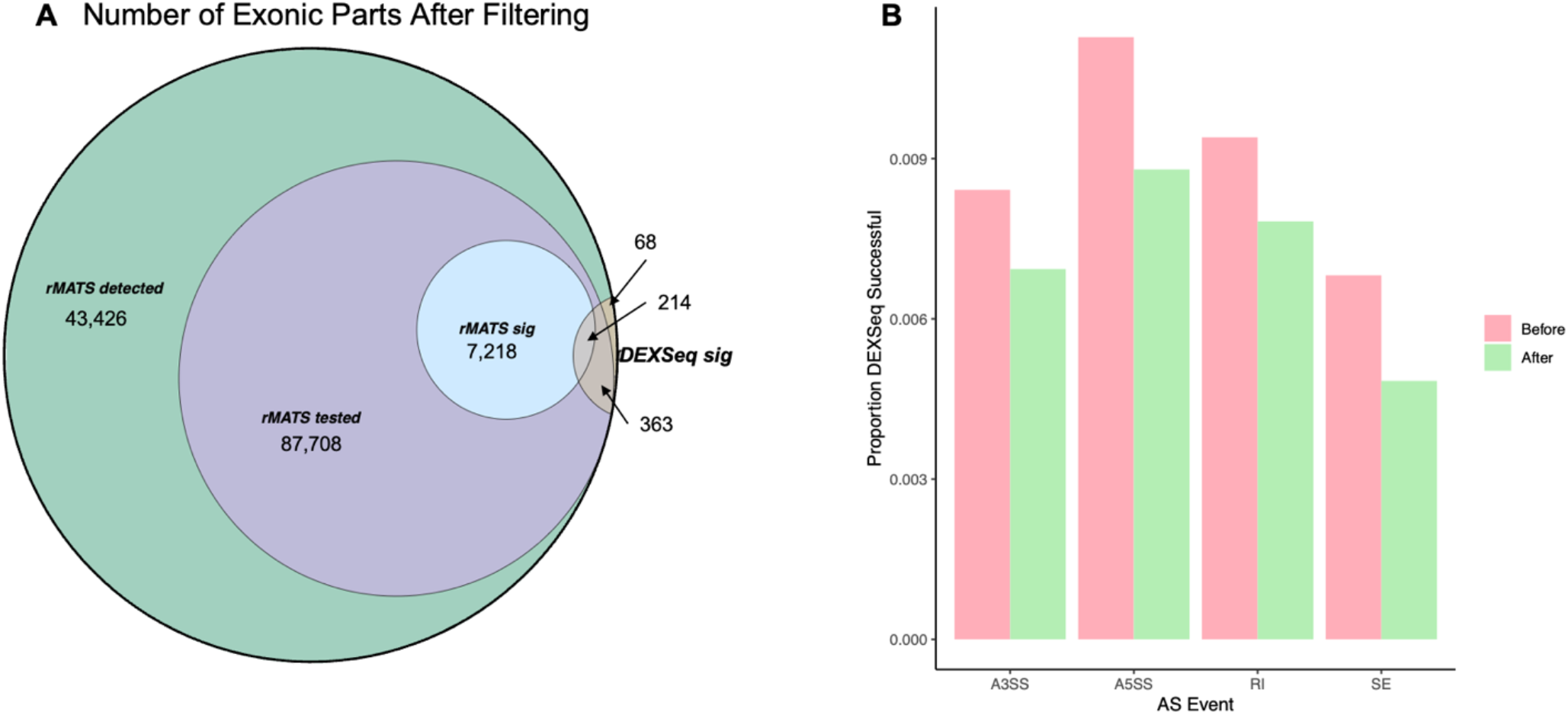
Direct comparison of AS events detected by DEXSeq before and after filtering for exons involved in AS. **A)** Venn Diagram categorizing DEXSeq exonic parts after filtering DEXSeq exons involved in AS. **B)** Barplot of significant exons before and after filtering for exons involved in AS. The pink bars represent the proportion of exons that DEXSeq detected as significant (adjusted p-value ≤ 0.05) over the total number of exons involved in the AS event before filtering the DEXSeq tested exons involved in AS. The green bars represent the proportion of exons that DEXSeq detected as significant over the total number of exons involved in the AS event after filtering the DEXSeq tested exons involved in AS.

### 3.6 Examination of the events missed by DEXSeq shows its lower sensitivity on exons that are part of multiple transcripts

We already know that the mis-identification of the exons described in section 3.3, and the multiple testing described in section 3.5 are contributing to the lower sensitivity observed in DEXSeq, but we wanted to examine the data further in relation to isoform complexity. DEXSeq fits a linear model that tests the effect of the interaction term condition:exon, where the exon is a factor with two levels: the number of reads mapping to the exonic part in question (level *this*), and the sum of the read counts from all other exonic parts of the same gene (level *others*). We formulated two hypotheses that could affect the sensitivity: first, the introduction of noise due to combining counts across many different isoforms into a summary count of “*others*”, especially if there are multiple isoforms differentially used simultaneously; second, the collapsing of signal when the exonic count is part of multiple isoforms, and only one of the isoforms is differentially used. To test these hypotheses, we looked at the relationship between the proportion of significant events detected vs. number of transcripts that are involved in “*this”* vs. “*others”*.

Fig 8A shows the proportion of significant events based on the number of isoforms that a gene has. Regardless of the number of isoforms, rMATS detects more significant AS events compared to DEXSeq. However, there was no change in the difference in proportion between rMATS and DEXSeq in relation to isoform complexity. Fig 8B shows the proportion of significant events based on the number of transcripts this exonic part is a member of. Again, we confirm that rMATS detects more significant events than DEXSeq. But, here, the overall proportion is even higher for rMATS than DEXSeq when the exonic part is a member of a large number of transcripts. We divided the transcript membership into two groups, less than 30 and more than 30. The difference in the proportion of significant events between rMATS and DEXSeq was significantly larger, when the exonic part was a member of more than 30 transcripts (*P*=0.04585). Given that this experiment is based on empirical data without ground truth, it is hard to conclude whether the lower detection seen in DEXSeq with higher transcript complexity is False Negatives for DEXSeq or False Positives for rMATS. At most, we can conclude that the isoform complexity surrounding the focal exon impacts the detection differently between the two approaches.

**Figure 8.**
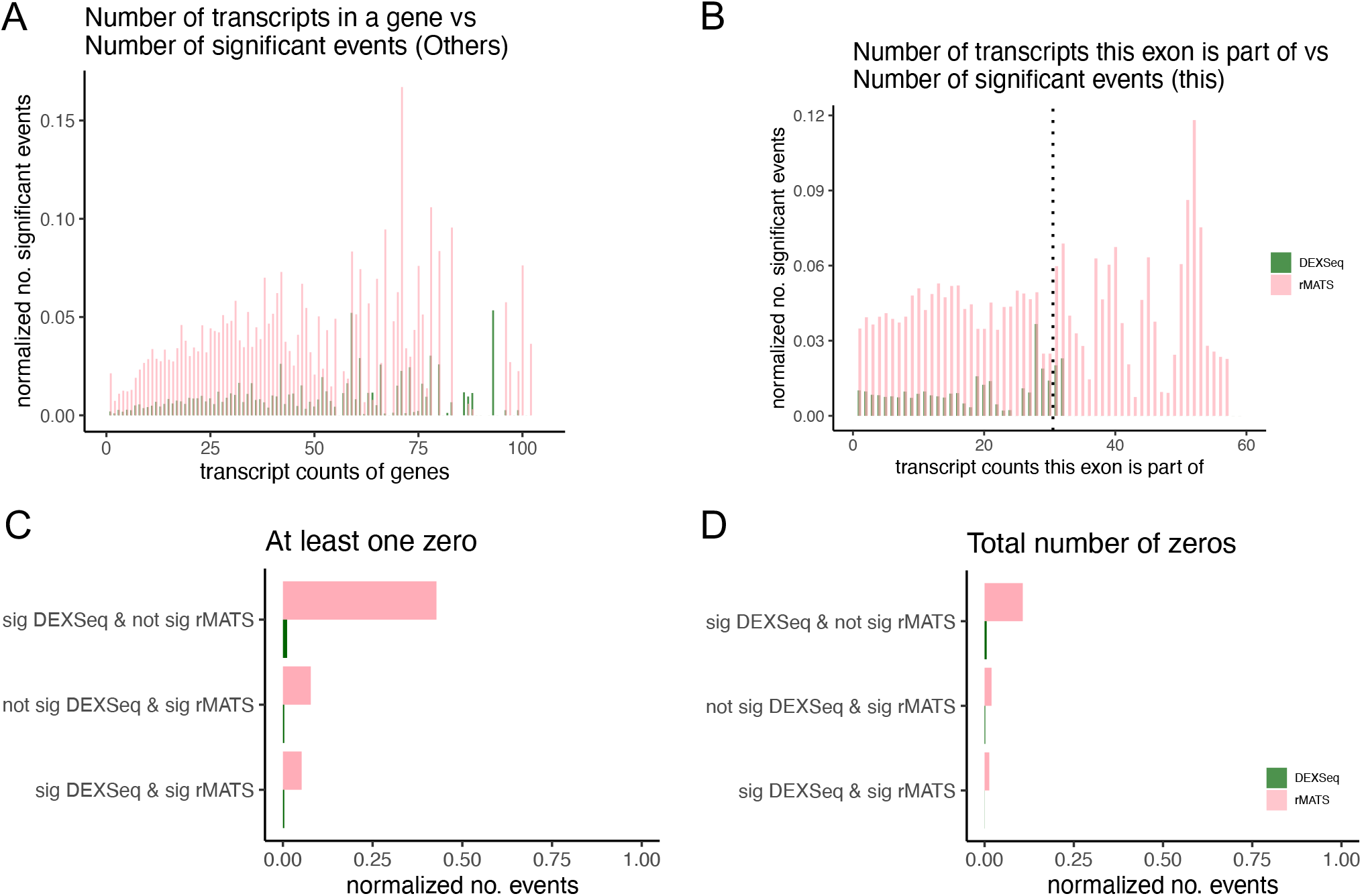
Detection by rMATS and DEXSeq in relation to isoform complexity and sparsity of read counts. A) Barplot showing the number of significant AS events (FDR ≤ 0.05) detected by rMATS (pink bars) and DEXSeq (dark green bars) based on the number of transcripts in a gene. B) Barplot showing the number of significant events (FDR ≤ 0.05) based on the number of transcripts an exon is part of. The dotted line represents the division between exons part of 30 or less transcripts vs exons part of more than 30 transcripts. C) Barplot showing the number of events that are significant (FDR ≤ 0.05) in DEXSeq and not significant (FDR > 0.05) in rMATS (top), not significant in DEXSeq and significant in rMATS (middle), and significant in DEXSeq and rMATS (bottom) based on the events having at least one zero sum read counts in B naive or CD8^+^ naive cells. D) Barplot showing the number of events based on the total number of zero sum read counts in B naive or CD8^+^ naive cells.

### 3.7 Examination of the events missed by rMATS shows its lower sensitivity on junctions that have low read coverage

In turn, we investigated the events that are missed by rMATS but identified by DEXSeq to understand the weakness of rMATS. One obvious difference is that exons involved in Alternative TSS and TTS will be detected by DEXSeq but missed by rMATS. This is likely the largest source of discrepancy. But, if we limit ourselves just to events that are true Alternative Splicing events, *i*.*e*. the events that are detected and tested by rMATS, we hypothesized that the failure to detect by rMATS may be due to the sparse read coverage at splice junctions. To test this hypothesis, we categorized all the AS events tested by rMATS into three categories: events detected by both software, events detected by DEXSeq but missed by rMATS, and events detected by rMATS but missed by DEXSeq. We then examined how many events had zero read counts when summed across all five samples, which we termed zero-sum cases. Figure 8C shows the number of events with at least one zero-sum count among the four counts (B cell inclusion, B cell exclusion, T cell inclusion, T cell exclusion) for rMATS, and the number of exonic parts with at least one zero-sum count among the two counts (B cell exonic part, T cell exonic part) for DEXSeq. As expected, there were more zero-sum cases among inclusion and exclusion counts in rMATS compared to exonic part counts in DEXSeq. Nevertheless, there was an even stronger excess of zero-sum cases among events called significant by DEXSeq but missed by rMATS, with 48% of these events having at least one zero-sum count. In contrast, among events there were called significant by rMATS but missed by DEXSeq, only 0.37% of exonic parts had at least one zero-sum count. This disparity underscores the vulnerability of rMATS to the sparsity of reads at splice junctions, a limitation not observed in DEXSeq (Figure 8C). We also examined at the proportion of zero-sum counts among all possible counts and found a parallel trend (Figure 8D).

### 3.8 Integrating the results from the two approaches produces a comprehensive list of AS detected between B cells and T cells

Based on the mapping between the events and exonic parts, we were able to integrate the results from DEXSeq and rMATS. This produced a comprehensive list of AS events and exons that are differentially used between purified B cells and T cells, including the events and exons that are detected by both software and are high confidence events (Figure 9) This list included some genes that were previously reported to express various isoforms in B cell or T cells, including CD37 [23], PIK3R1 [24], IRF7 [25], and several other genes that have been reported in mice (NCF1, NOSIP, TPP1, ZFAND1, etc) [26]. We expected to identify common genes that are differentially spliced in the comparison of B cell vs. T cells and the comparison of Raji vs. Jurkat cell lines. But, the list were largely different. The common genes that were found between the two comparison were ERAP2 and WARS1. There are several genes with functions in DNA repair that were uniquely differentially spliced in the Raji vs. Jurkat comparison, that included *ERCC1, FANCC, FANCA*, and *FNA1* (Supplementary Figure S14).

**Figure 9.**
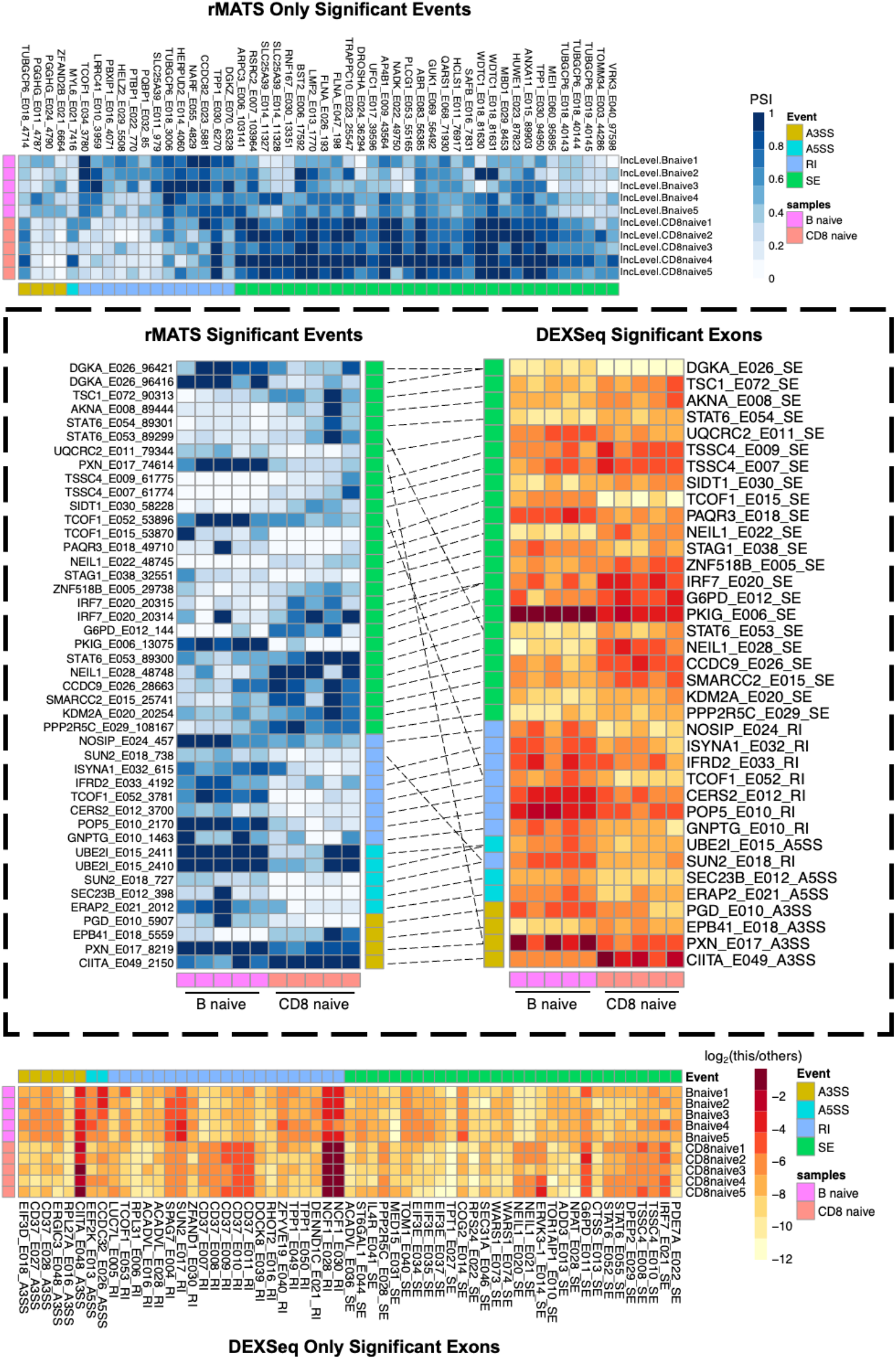
Heatmaps of significant genes detected by rMATS and DEXSeq on B naive and CD8^+^ naive cells. The columns of the heatmap represent the genes grouped by the four AS events: khaki = alternative 3’ splice site (A3SS), turquoise = alternative 5’ splice site (A5SS), blue = retained intron (RI), green = skipped exon (SE). The rows represent the samples: violet = B naive cells, salmon = CD8^+^ naive cells. The top heatmap display the inclusion levels (PSI) of the significant genes (FDR ≤ 0.05) only detected by rMATS. Annotated on the columns are the gene symbols followed by the DEXSeq exon and then the unique rMATS ID. The bottom heatmap display the this/others counts log2 transformed of the significant genes (adjusted p-value ≤ 0.05) only detected by DEXSeq. The gene symbols are followed by the DEXSeq exon and then the corresponding AS event. The middle heatmaps represent significant genes that are shared between rMATS (left) and DEXSeq (right) connected by dashed lines.

## 4 Discussion

In this study, we introduced a novel splicing graph, GrASE, that offers a means to directly compare splice junction-based and exon-based methods at both the event and exon levels. This novel approach enabled the precise localization of discrepancies between the two approaches, providing insights into their respective strengths and limitations.

DEXSeq, despite its name suggesting a focus on exons, occupies an intermediate space between exon-level and isoform-level detection. It is capable of detecting differential usage due to Alternative Splicing, but also Alternative Transcription Start Site (TSS) or Alternative Transcription Termination Site (TTS) events. Designed to test interaction between the experimental condition and *this vs. others* count levels, we observed that the software will frequently detect the interaction driven by changes in “*others*” rather than “*this*”, especially when there are large changes in “*others*” due to the differences in overall length of the isoforms that are undergoing switches due to alternative TSS or TTS. This characteristic of the model has been clearly described in previous publications by the authors [21,22], but this nature of the model, or how much “*others*” can differ depending on the isoforms in empirical data may not be well understood or appreciated by the users of the software. At the gene level, DEXSeq typically performs as expected, detecting changes in the isoform usage that manifests at some of the exons. But in some cases, the model mis-identified the focal exons that were involved in the isoform switch, because there is nothing in the design to prioritize change in “*this”* over “*others”*. This resulted in unpredictability in the detection of the focal exon, and discrepancies between the results of event-based *vs*. exon-based detection.

On the other hand, the strengths of DEXSeq and the weaknesses of rMATS became apparent where it exhibited higher specificity in scenarios with larger variance. Although the simulation only considered dispersion as small as 1/30 of the transcript’s mean, further reduction of dispersion would likely have highlighted DEXSeq’s capabilities even more prominently. Additionally, we observed the weaknesses of rMATS when there were a small number of reads mapping to junctions as expected. This was evidenced by the prevalence of zero-sum counts among the events missed by rMATS but detected by DEXSeq. It was also seen in the incorrect estimation of PSI level change when the exclusion counts were zero.

One limitation of this study is that we relied on the detection of events by rMATS as the starting point of our mapping between events and exons. rMATS limits their detection of AS to simple binary events of specific types. An alternative solution to this problem is to catalog splicing bubbles [18] or local splicing variants (LSV) [8,27] from the annotations as the starting point for the mapping. Looking at bubbles or LSVs will theoretically give us a comprehensive catalog of events, not limited by the complexity or the type of event. Unfortunately, with bubbles, it quickly becomes difficult to identify the exon to event mappings between nested bubbles in the annotation structure. Note that the algorithm for mapping between event and focal exonic part described in section 2.1 only works for bubbles with binary alternative paths. Where there are more than two paths possible, we must determine which two paths to compare to define the focal exonic part, and it can quickly become combinatorically complex. Nested bubbles can lead to uncertainty on which event is responsible for the exon usage change. On the other hand, with LSVs, we give up on the information connecting the source and sink that defines the event, and aim for a modest resolution at either of the junctions rather than capturing the full event. This is an inherent limitation of the short reads that will only be solved with integration of the long read sequencing data.

In summary, we have proposed a method for bridging two methodologies in AS analysis. Through performance benchmarking, we identified strengths and weaknesses in each approach. The large discrepancy observed between the two most popular software used in the field can be understood as the large difference in the models of what they are designed to detect. It also underscores the fact that AS detection based on short read sequencing remains a challenging problem that is far from being solved. The weaknesses identified through this comparison highlights areas for improvement, and will aid in improving the detection of AS utilizing short reads.

## References

1. Su C-H, D D, Tarn W-Y. Alternative Splicing in Neurogenesis and Brain Development. Front Mol Biosci 2018;5.

2. Liu H, He L, Tang L. Alternative Splicing Regulation and Cell Lineage Differentiation. Curr Stem Cell Res Ther 2012;7:400–6.

3. Evsyukova I, Somarelli JA, Gregory SG et al. Alternative splicing in multiple sclerosis and other autoimmune diseases. RNA Biol 2010;7:462–73.

4. Li H, Wang Z, Ma T et al. Alternative splicing in aging and age-related diseases. Transl Med Aging 2017;1:32–40.

5. Bergsma AJ, van der Wal E, Broeders M et al. Chapter Three - Alternative Splicing in Genetic Diseases: Improved Diagnosis and Novel Treatment Options. In: Loos F (ed.). International Review of Cell and Molecular Biology. Vol 335. Academic Press, 2018, 85–141.

6. Shen S, Park JW, Lu Z et al. rMATS: Robust and flexible detection of differential alternative splicing from replicate RNA-Seq data. Proc Natl Acad Sci 2014;111:E5593–601.

7. Trincado JL, Entizne JC, Hysenaj G et al. SUPPA2: fast, accurate, and uncertainty-aware differential splicing analysis across multiple conditions. Genome Biol 2018;19:40.

8. Vaquero-Garcia J, Barrera A, Gazzara MR et al. A new view of transcriptome complexity and regulation through the lens of local splicing variations. Valcárcel J (ed.). eLife 2016;5:e11752.

9. Anders S, Reyes A, Huber W. Detecting differential usage of exons from RNA-Seq data. Nat Preced 2012:1–1.

10. Hartley SW, Mullikin JC. Detection and visualization of differential splicing in RNA-Seq data with JunctionSeq. Nucleic Acids Res 2016;44:e127.

11. Soneson C, Matthes KL, Nowicka M et al. Isoform prefiltering improves performance of count-based methods for analysis of differential transcript usage. Genome Biol 2016;17:12.

12. Jiang M, Zhang S, Yin H et al. A comprehensive benchmarking of differential splicing tools for RNA-seq analysis at the event level. Brief Bioinform 2023;24:bbad121.

13. Olofsson D, Preußner M, Kowar A et al. One pipeline to predict them all? On the prediction of alternative splicing from RNA-Seq data. Biochem Biophys Res Commun 2023;653:31–7.

14. Ding L, Rath E, Bai Y. Comparison of Alternative Splicing Junction Detection Tools Using RNA-Seq Data. Curr Genomics 2017;18:268–77.

15. Liu R, Loraine AE, Dickerson JA. Comparisons of computational methods for differential alternative splicing detection using RNA-seq in plant systems. BMC Bioinformatics 2014;15:364.

16. Mehmood A, Laiho A, Venäläinen MS et al. Systematic evaluation of differential splicing tools for RNA-seq studies. Brief Bioinform 2020;21:2052–65.

17. Sterne-Weiler T, Weatheritt RJ, Best AJ et al. Efficient and Accurate Quantitative Profiling of Alternative Splicing Patterns of Any Complexity on a Laptop. Mol Cell 2018;72:187-200.e6.

18. Sammeth M. Complete Alternative Splicing Events Are Bubbles in Splicing Graphs. J Comput Biol 2009;16:1117–40.

19. Bindreither D, Carlson M, Morgan M et al. SplicingGraphs: Create, manipulate, visualize splicing graphs, and assign RNA-seq reads to them. R Package Version 1380 2022.

20. Frazee AC, Jaffe AE, Langmead B et al. Polyester: simulating RNA-seq datasets with differential transcript expression. Bioinformatics 2015;31:2778–84.

21. Anders S, Reyes A, Huber W. Detecting differential usage of exons from RNA-seq data. Genome Res 2012;22:2008–17.

22. Reyes A, Huber W. Alternative start and termination sites of transcription drive most transcript isoform differences across human tissues. Nucleic Acids Res 2018;46:582–92.

23. Byrne A, Beaudin AE, Olsen HE et al. Nanopore long-read RNAseq reveals widespread transcriptional variation among the surface receptors of individual B cells. Nat Commun 2017;8:16027.

24. Fruman DA, Snapper SB, Yballe CM et al. Impaired B Cell Development and Proliferation in Absence of Phosphoinositide 3-Kinase p85α. Science 1999;283:393–7.

25. Ip JY, Tong A, Pan Q et al. Global analysis of alternative splicing during T-cell activation. RNA N Y N 2007;13:563–72.

26. Ergun A, Doran G, Costello JC et al. Differential splicing across immune system lineages. Proc Natl Acad Sci 2013;110:14324–9.

27. Vaquero-Garcia J, Aicher JK, Jewell S et al. RNA splicing analysis using heterogeneous and large RNA-seq datasets. Nat Commun 2023;14:1230.

