## Supplementary Information for "A novel splicing graph allows a direct comparison between exon-based and splice junction-based approaches to alternative splicing detection"

### 1 Supplementary Tables

**Supplementary Table S1**: Number of simulations for each fold change

| FC | Simulations |
|---|---|
| 0 | 36 |
| +1 | 18 |
| -1 | 18 |
| +2 | 8 |
| -2 | 8 |
| +4 | 4 |
| -4 | 4 |
| +8 | 2 |
| -8 | 2 |

FC = log2 fold change

**Supplementary Table S2**: Number of simulations for each read depth

| Read Depth | Simulations |
|---|---|
| 1 | 250 |
| 2 | 125 |
| 3 | 60 |
| 4 | 30 |
| 5 | 15 |
| 6 | 8 |
| 7 | 4 |
| 8 | 2 |
| 9 | 1 |
| 10 | 1 |
| 50 | 1 |
| 100 | 1 |

| | *this* | | *others* | | *p-values* | |
|---|---|---|---|---|---|---|
| Skipped Exon | control | case | control | case | raw *p-value* | adjusted *p-value* |
| ENSG00000081181.8:E006 | 499 | 2233 | 4242 | 18249 | 0.148 | 0.155 |
| ENSG00000163131.11:E018 | 487 | 2055 | 10320 | 41949 | 0.130 | 0.137 |
| ENSG00000186710.11:E007 | 563 | 2724 | 3320 | 16405 | 0.357 | 0.363 |
| ENSG00000116014.10:E004 | 429 | 2128 | 2936 | 15093 | 0.158 | 0.165 |
| ENSG00000102076.10:E006 | 500 | 2520 | 2740 | 14314 | 0.132 | 0.139 |
| ENSG00000121481.11:E008 | 552 | 2378 | 5071 | 22641 | 0.114 | 0.121 |
| ENSG00000117222.14:E015 (RBBP5) | 743 | 3999 | 43138 | 204339 | 0.091 | 0.787 |

Supplementary Table S3. Simulated positive skipped exon cases that DEXSeq failed to detect. Counts from *this* and *others* columns were averaged for the five samples in control and case.

#### 2 Supplementary Figures

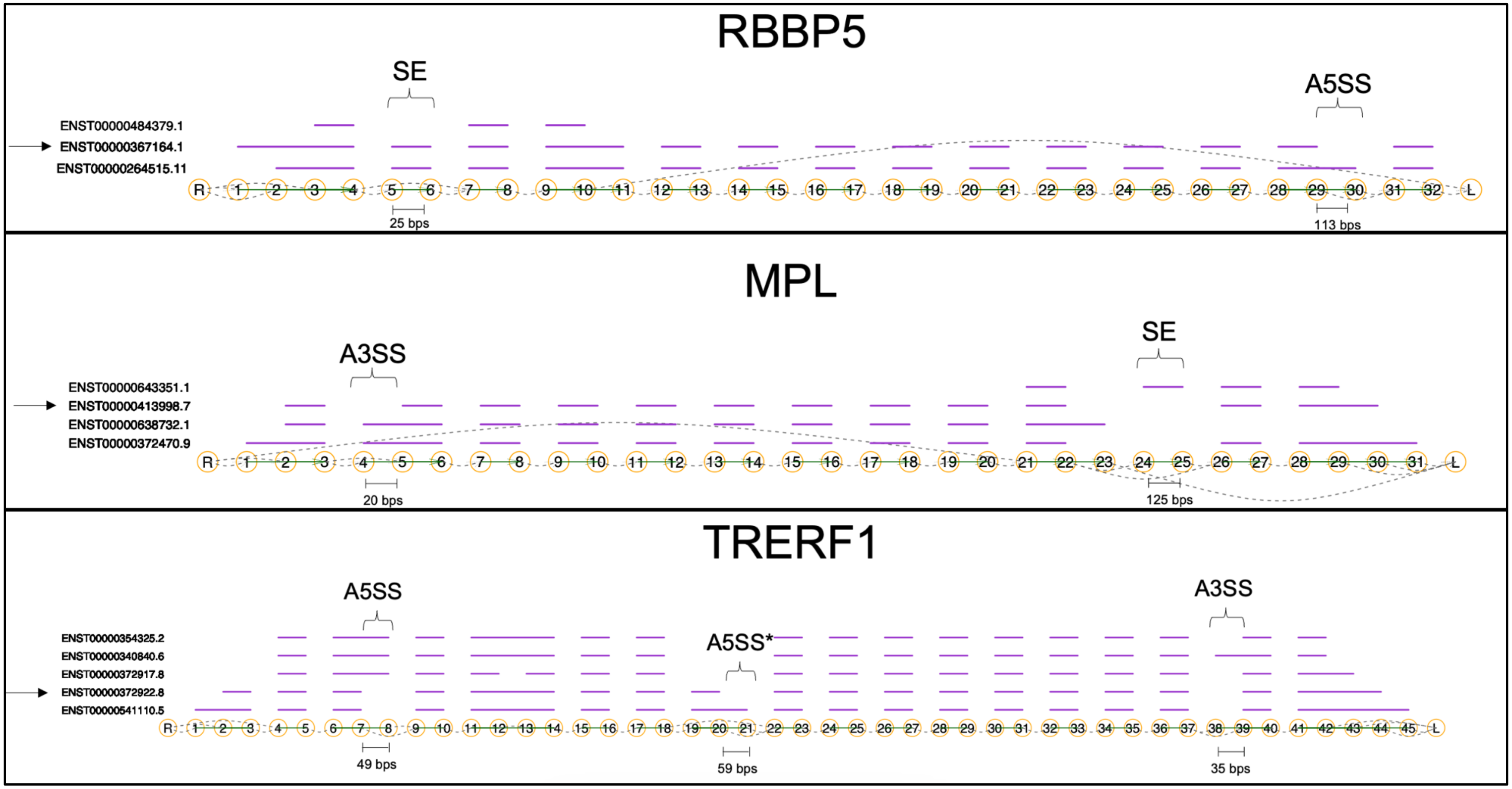


Supplementary Figure S1: Splicing graphs of RBBP5, MPL and TRERF1. Exons that were used for the simulations are annotated with their associated splicing event and exon length in base pairs (bps). The arrows adjacent to the transcript names are the transcripts we manipulated in our simulations. SE = Skipped Exon; A5SS = Alternative 5' Splice Site; A3SS = Alternative 3' Splice Site; A5SS* = Alternative 5' Splice Site used for simulation.

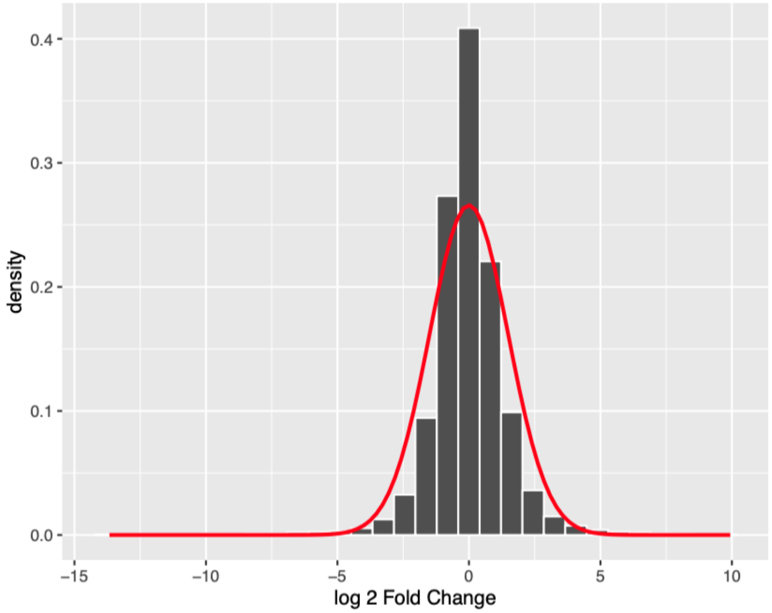


**Supplementary Figure S2**: Histogram showing the distribution of the log2 fold change resulting from the differential gene expression analysis performed using DESeq2 between B naive and $CD8^+$ naive cells. The distribution was fitted using a normal distribution curve with a mean of -0.003 and a standard deviation of 1.5.

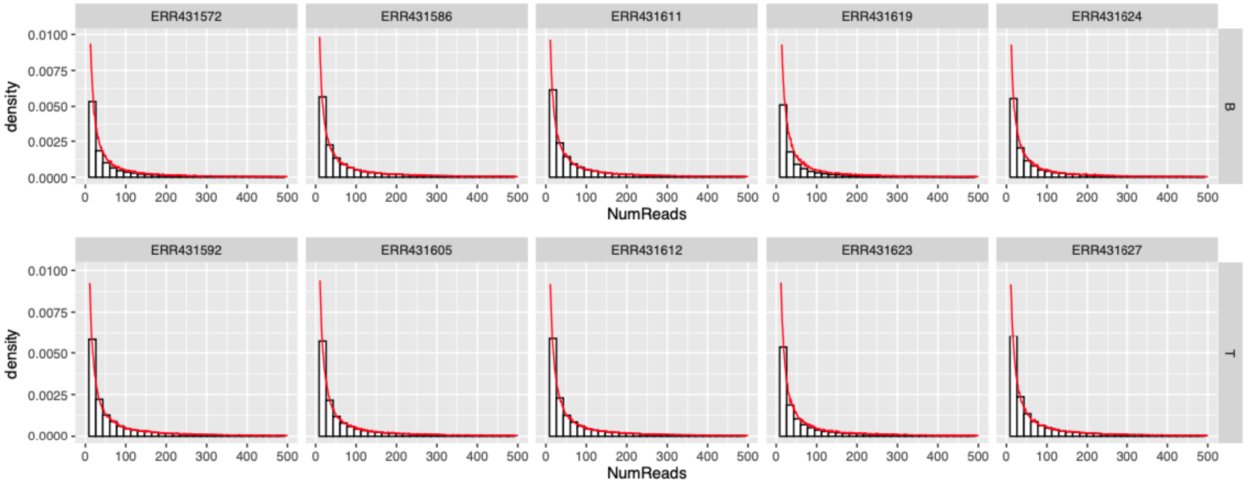


**Supplementary Figure S3**: Histogram showing the distribution of the read counts mapped to the transcripts from paired-end bulk RNA-seq reads of the five B naive cells (top row) and the five $CD8^+$ naive cells (bottom row). The distributions had an average decay rate of 0.024.

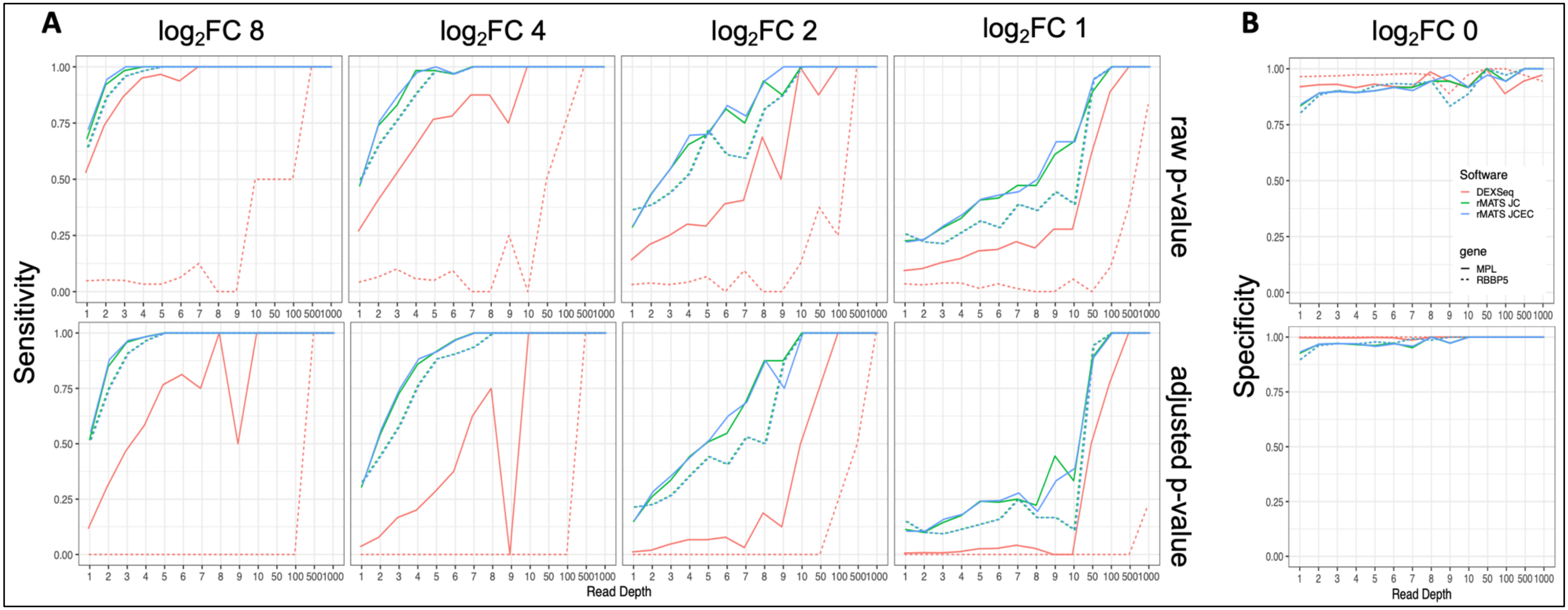


Supplementary Figure S4: Sensitivity and specificity of MPL and RBBP5 skipped exon (SE) simulations at dispersion (σ) ≈ 0.33. A) Sensitivity results of the SE event in MPL (solid line) RBBP5 (dotted line). Each line plot represents the sensitivity (y-axis) vs the read depth (x-axis) for different log2 fold changes (8,4,2,1). The green line represents the performance based on rMATS's junction counts (JC), the blue line is rMATS's junction and exon counts (JCEC), and the red line is DEXSeq. The top panel represents the sensitivity based on the raw p-value of the test, while the bottom panel represents the adjusted p-value. B) Specificity results (log2 FC 0) of the SE event in MPL and RBBP5. The top panel represents the specificity based on the raw p-value of the test, while the bottom panel represents the adjusted p-value.

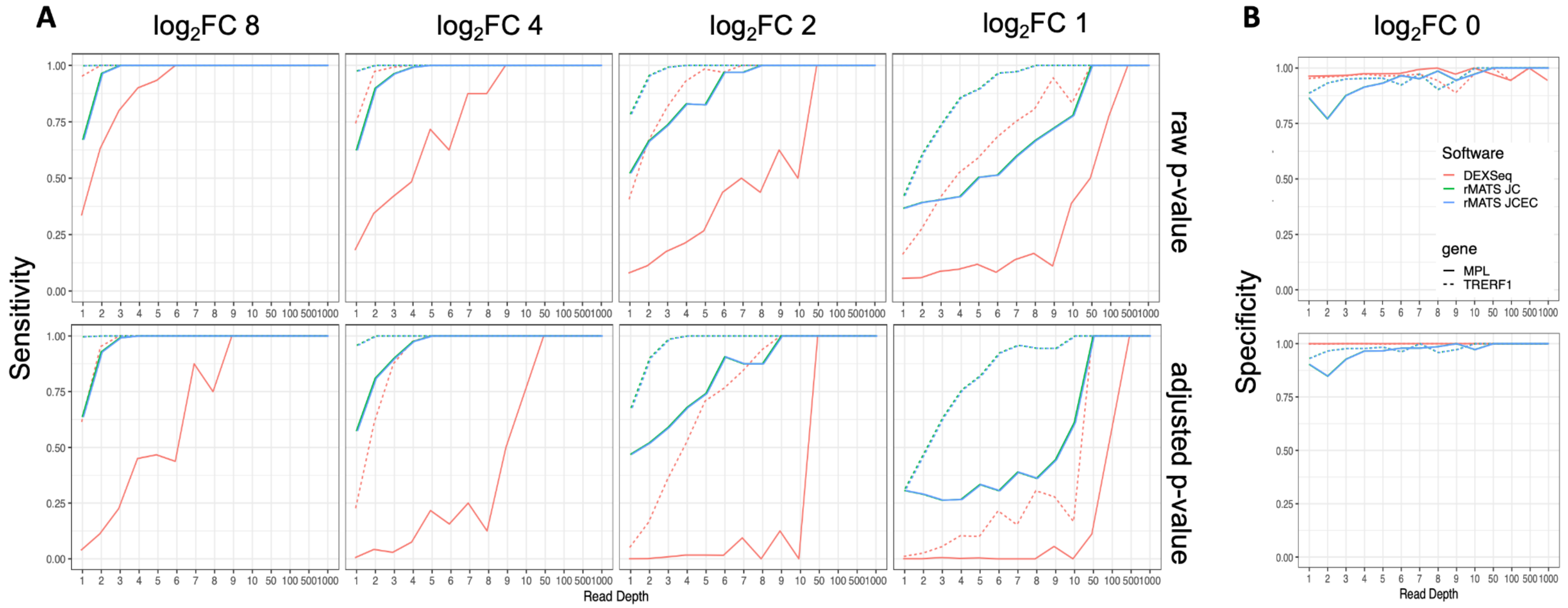


Supplementary Figure S5: Sensitivity and specificity of MPL and TRERF1 alternative 3' splice site (A3SS) simulations at dispersion (σ) ≈ 0.33. **A)** Sensitivity results of the A3SS event in MPL (solid line) TRERF1 (dotted line). Each line plot represents the sensitivity (y-axis) vs the read depth (x-axis) for different $\log_2$ fold changes (8,4,2,1). The green line represents the performance based on rMATS's junction counts (JC), the blue line is rMATS's junction and exon counts (JCEC), and the red line is DEXSeq. The top panel represents the sensitivity based on the raw p-value of the test, while the bottom panel represents the adjusted p-value. **B)** Specificity results ($\log_2$ FC 0) of the A3SS event in MPL and TRERF1. The top panel represents the specificity based on the raw p-value of the test, while the bottom panel represents the adjusted p-value.

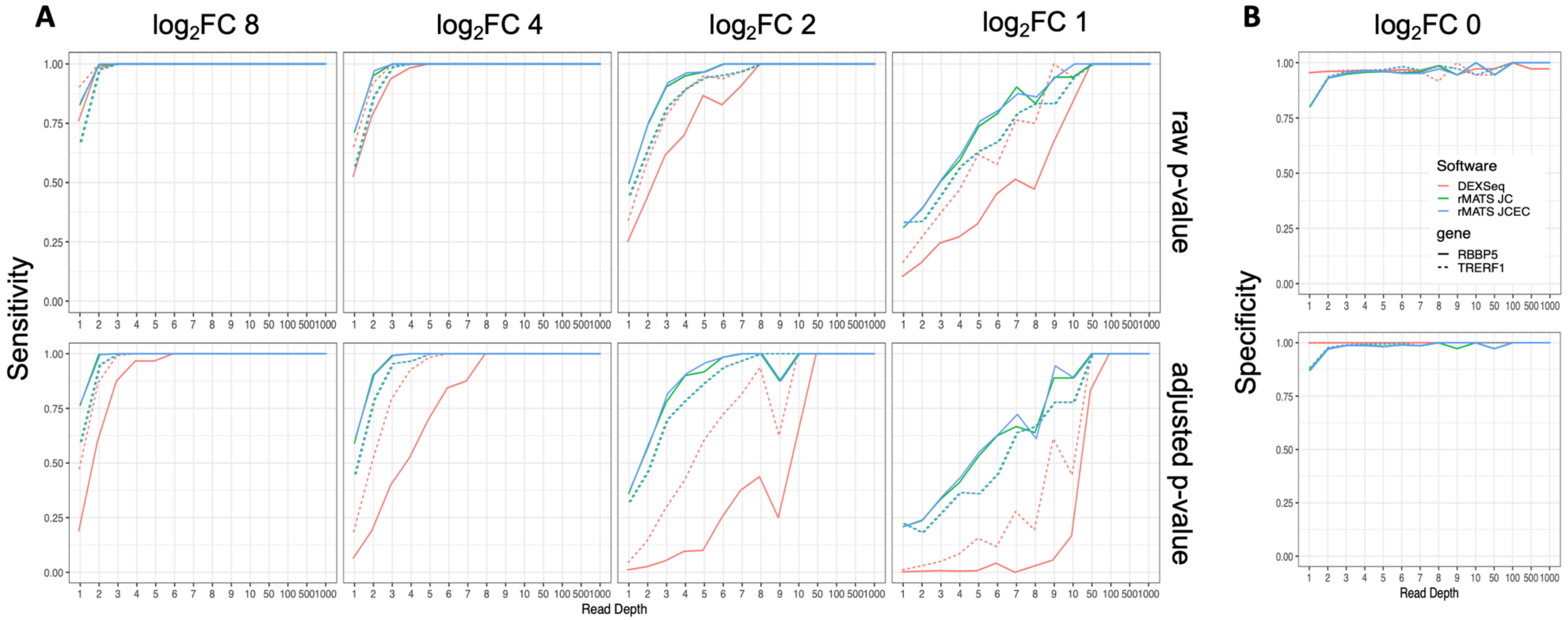


Suppelmentary Figure S6: Sensitivity and specificity of RBBP5 and TRERF1 alternative 5' splice site (A5SS) simulations at dispersion (σ) ≈ 0.33. A) Sensitivity results of the A5SS event in MPL (solid line) TRERF1 (dotted line). Each line plot represents the sensitivity (y-axis) vs the read depth (x-axis) for different log2 fold changes (8,4,2,1). The green line represents the performance based on rMATS's junction counts (JC), the blue line is rMATS's junction and exon counts (JCEC), and the red line is DEXSeq. The top panel represents the sensitivity based on the raw p-value of the test, while the bottom panel represents the adjusted p-value. B) Specificity results (log2 FC 0) of the A5SS event in RBBP5 and TRERF1. The top panel represents the specificity based on the raw p-value of the test, while the bottom panel represents the adjusted p-value.

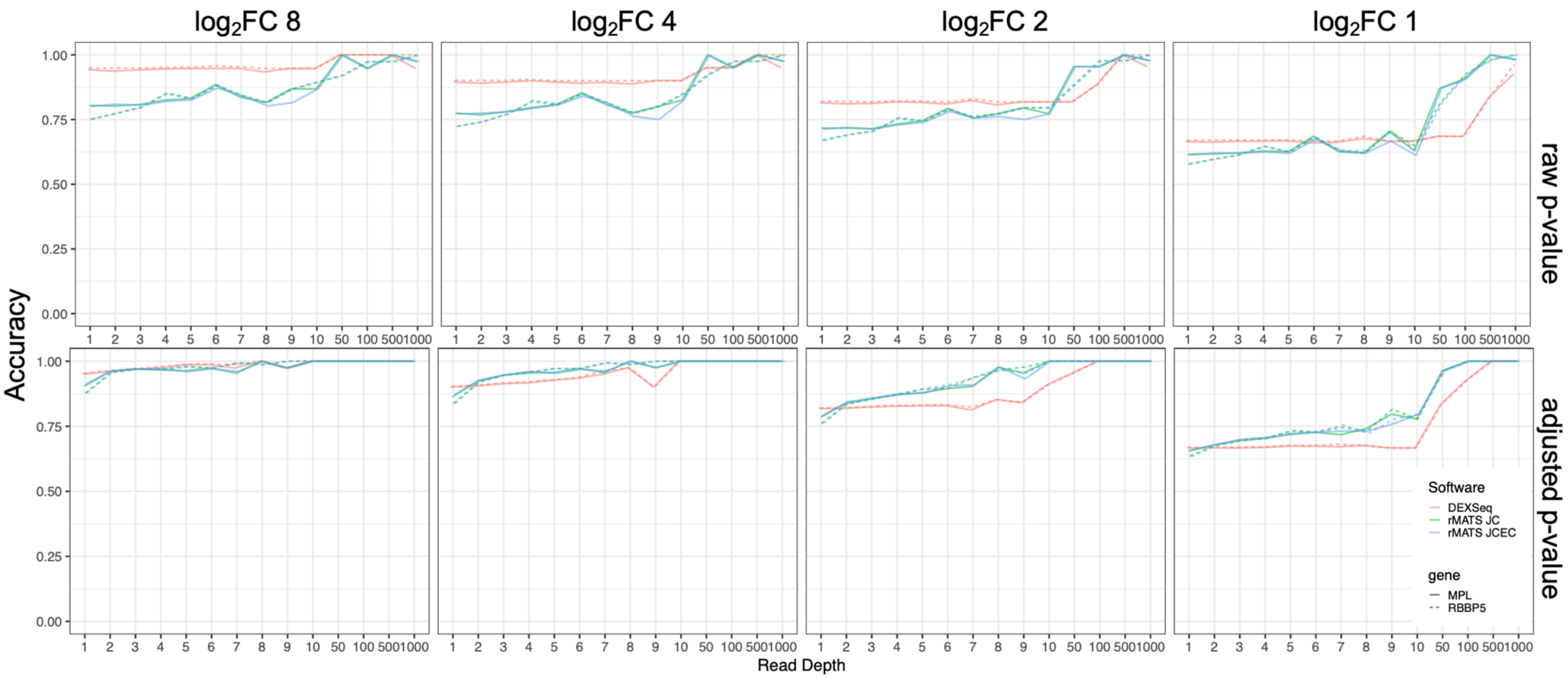


Supplementary Figure S7: Accuracy of MPL and RBBP5 skipped exon (SE) simulations at dispersion (σ) ≈ 0.033. Solid lines represent MPL and dotted lines are RBBP5. Each line plot represents the accuracy (y-axis) vs the read depth (x-axis) for different $\log_2$ fold changes (1, 2, 4 and 8). The green line represents rMATS's junction counts (JC), the blue line is rMATS's junction and exon counts (JCEC), and the red line is DEXSeq. The top panel represents the accuracy based on the raw p-value of the test, while the bottom panel represents the adjusted p-value.

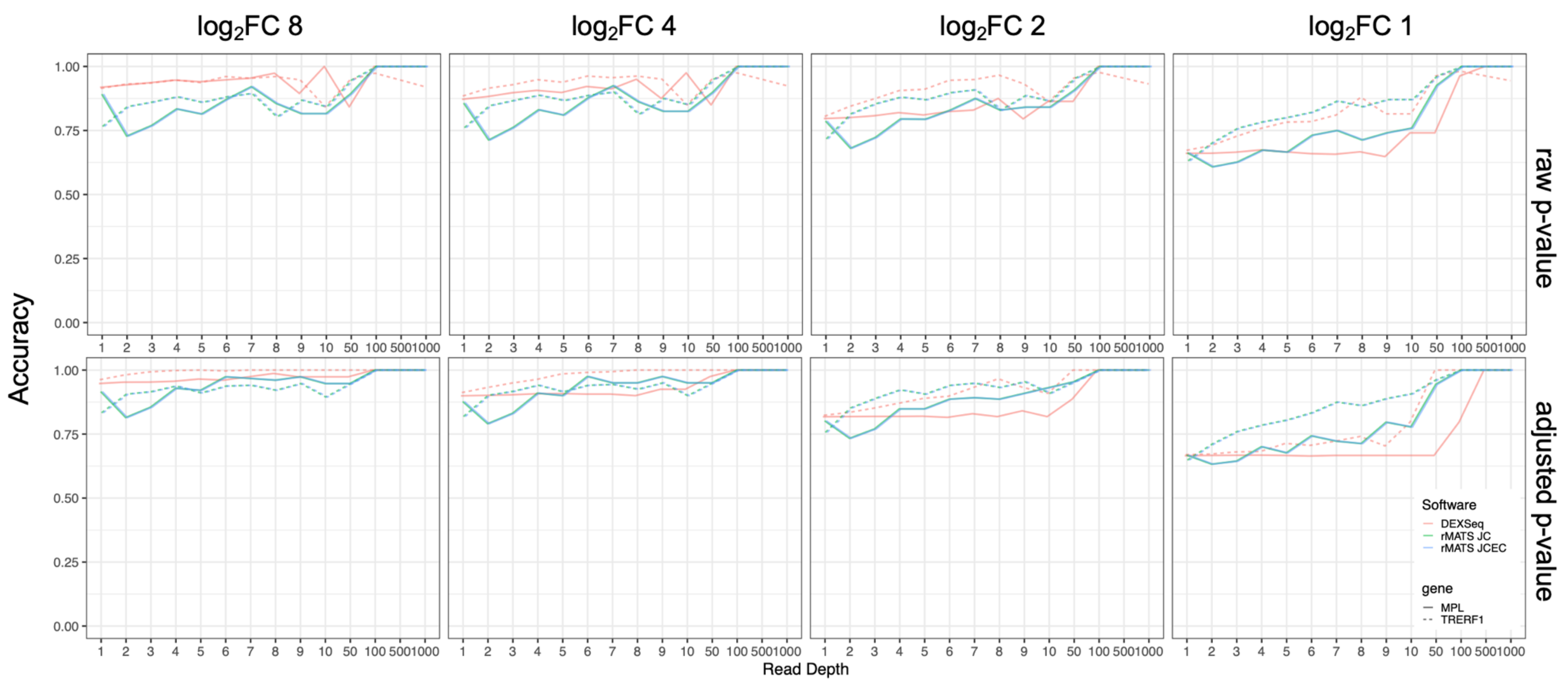


Supplementary Figure S8: Accuracy of MPL and TRERF1 alternative 3' splice site (A3SS) simulations at dispersion (σ) ≈ 0.033 . Solid lines represent MPL and dotted lines are TRERF1. Each line plot represents the accuracy (y-axis) vs the read depth (x-axis) for different $\log_2$ fold changes (1, 2, 4 and 8). The green line represents rMATS's junction counts (JC), the blue line is rMATS's junction and exon counts (JCEC), and the red line is DEXSeq. The top panel represents the accuracy based on the raw p-value of the test, while the bottom panel represents the adjusted p-value.

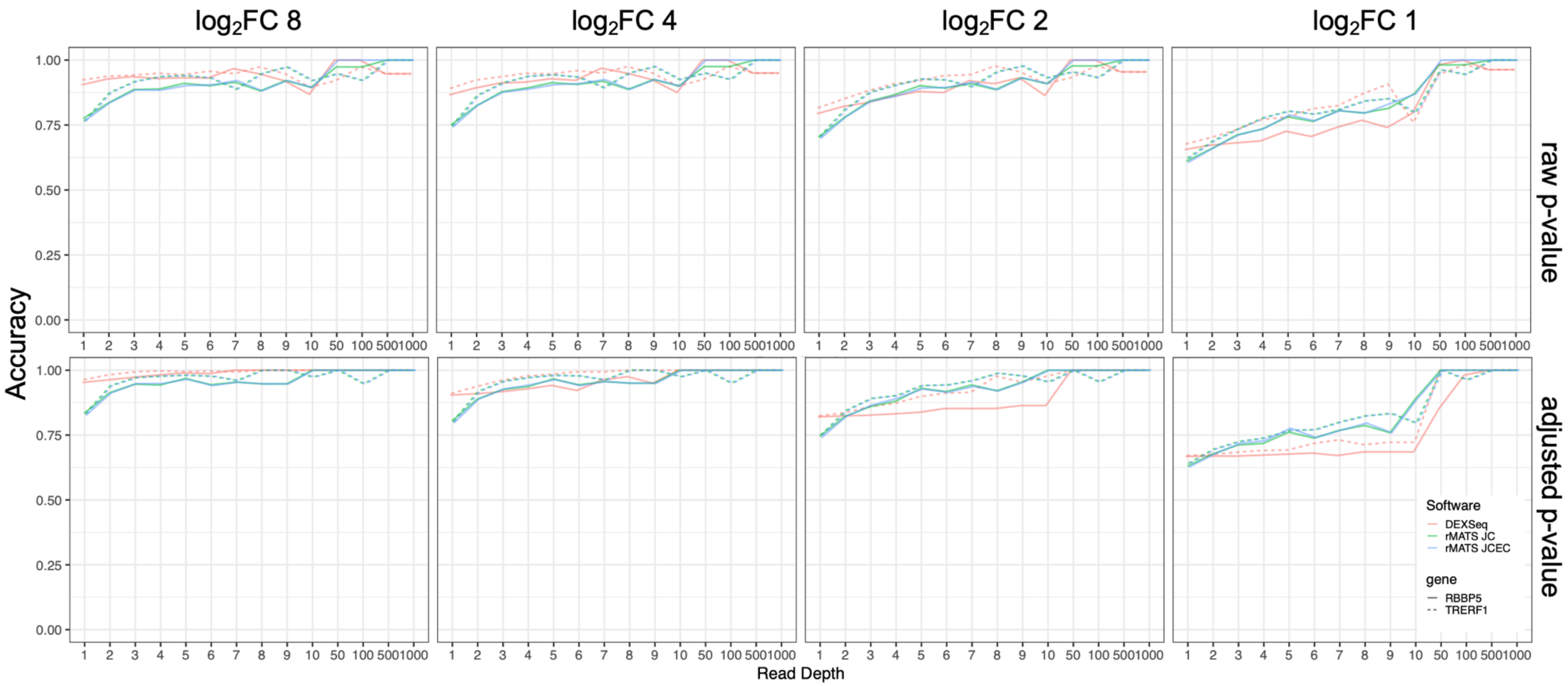


Supplementary Figure S9: Accuracy of RBBP5 and TRERF1 alternative 5' splice site (A5SS) simulations at dispersion (σ) ≈ 0.033. Solid lines represent RBBP5 and dotted lines are TRERF1. Each line plot represents the accuracy (y-axis) vs the read depth (x-axis) for different $\log_2$ fold changes (1, 2, 4 and 8). The green line represents rMATS's junction counts (JC), the blue line is rMATS's junction and exon counts (JCEC), and the red line is DEXSeq. The top panel represents the accuracy based on the raw p-value of the test, while the bottom panel represents the adjusted p-value.

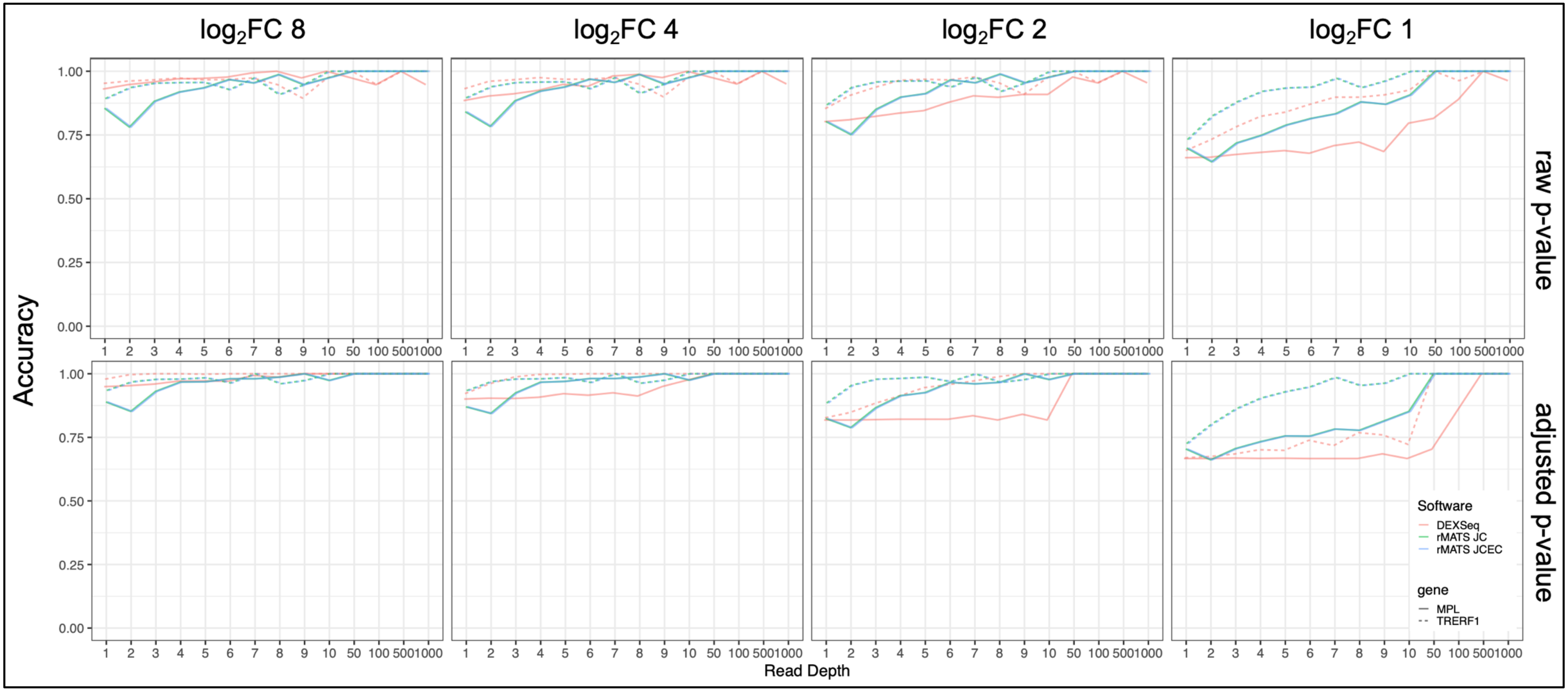


Supplementary Figure S10: Accuracy of MPL and TRERF1 alternative 3' splice site (A3SS) simulations at dispersion (σ) ≈ 0.33 . Solid lines represent MPL and dotted lines are TRERF1. Each line plot represents the accuracy (y-axis) vs the read depth (x-axis) for different $\log_2$ fold changes (1, 2, 4 and 8). The green line represents rMATS's junction counts (JC), the blue line is rMATS's junction and exon counts (JCEC), and the red line is DEXSeq. The top panel represents the accuracy based on the raw p-value of the test, while the bottom panel represents the adjusted p-value.

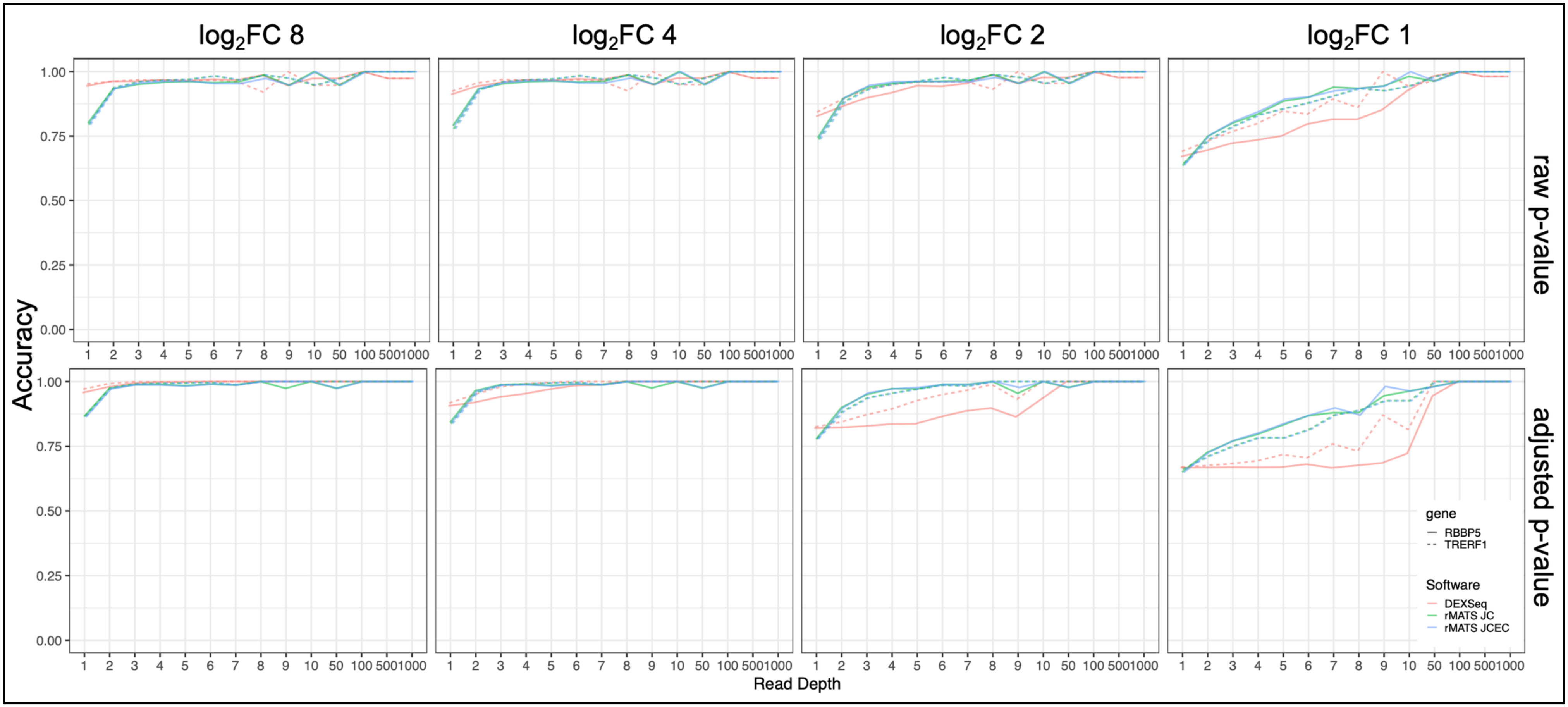


Supplementary Figure S11: Accuracy of RBBP5 and TRERF1 alternative 5' splice site (A5SS) simulations at dispersion (σ) ≈ 0.33. Solid lines represent RBBP5 and dotted lines are TRERF1. Each line plot represents the accuracy (y-axis) vs the read depth (x-axis) for different $\log_2$ fold changes (1, 2, 4 and 8). The green line represents rMATS's junction counts (JC), the blue line is rMATS's junction and exon counts (JCEC), and the red line is DEXSeq. The top panel represents the accuracy based on the raw p-value of the test, while the bottom panel represents the adjusted p-value.

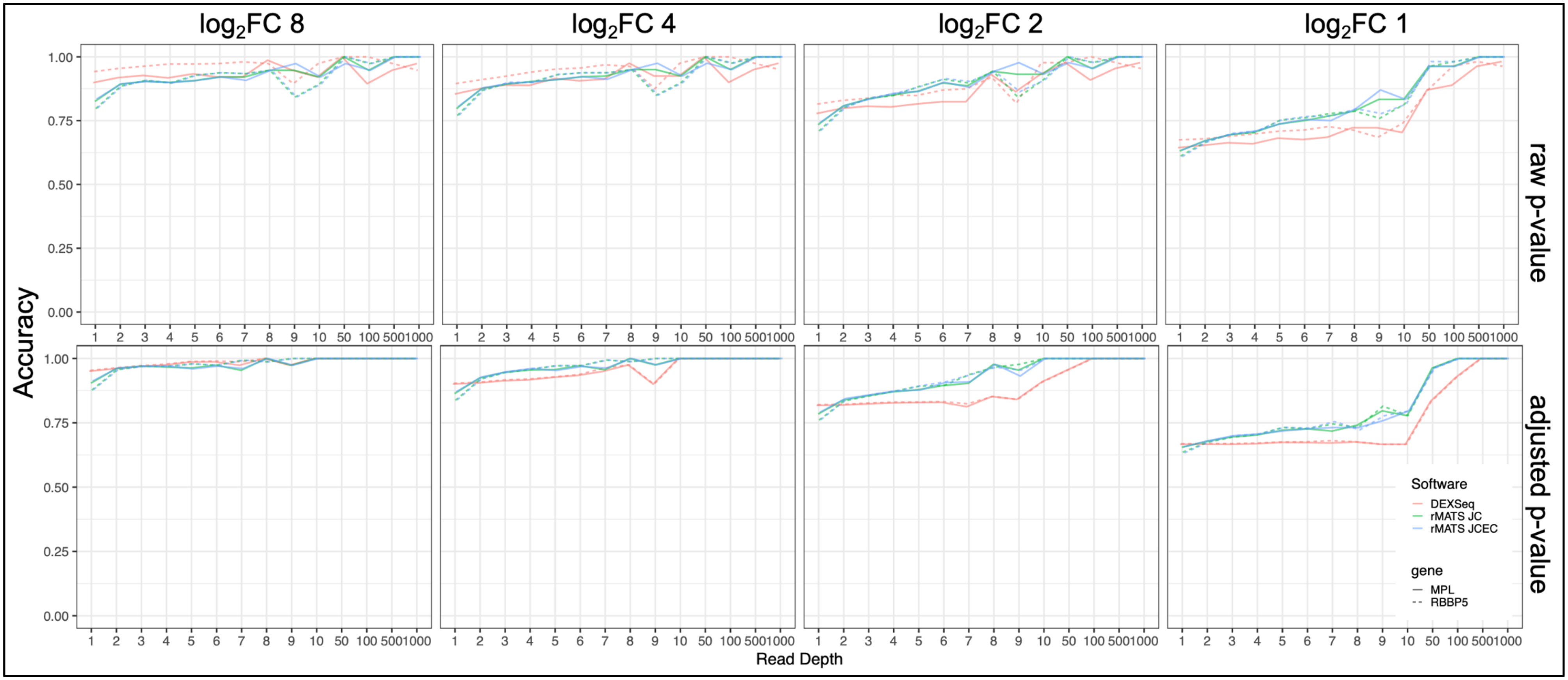


Supplementary Figure S12: Accuracy of MPL and RBBP5 skipped exon (SE) simulations at dispersion (σ) ≈ 0.33. Solid lines represent MPL and dotted lines are RBBP5. Each line plot represents the accuracy (y-axis) vs the read depth (x-axis) for different log2 fold changes (1, 2, 4 and 8). The green line represents rMATS's junction counts (JC), the blue line is rMATS's junction and exon counts (JCEC), and the red line is DEXSeq. The top panel represents the accuracy based on the raw p-value of the test, while the bottom panel represents the adjusted p-value.

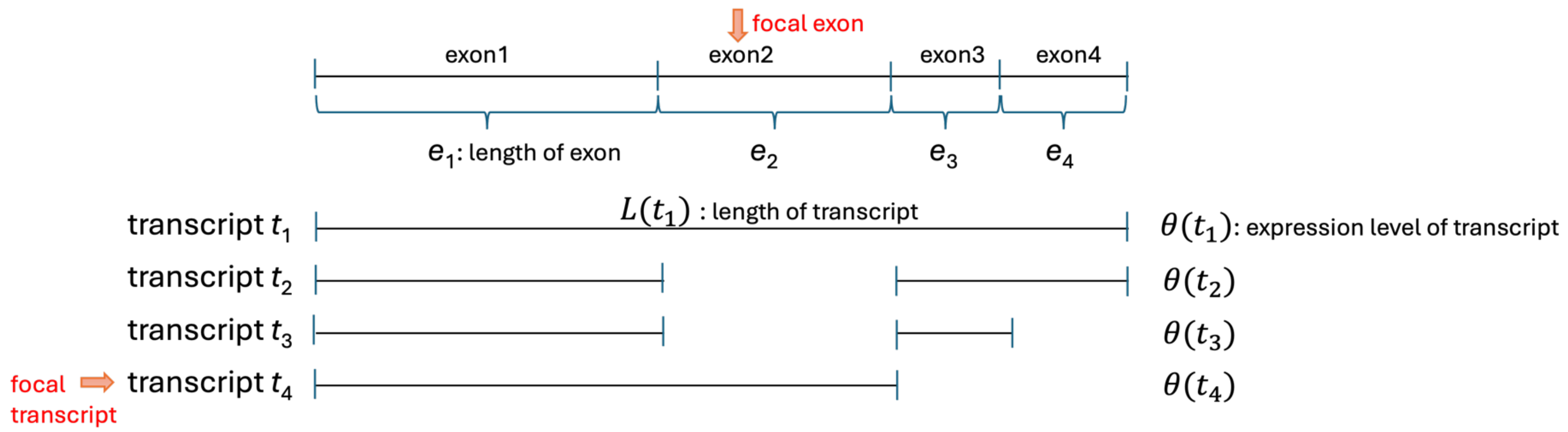


$$(lfc-1)\cdot\theta(t_f)\frac{e_f}{L(t_f)}\cdot\left[\sum_{t_i\in in}\frac{(L(t_f)-e_f)}{L(t_i)}\theta(t_i)-\sum_{t_i\in ex}\theta(t_i)-\sum_{t_i\in in}\frac{(L(t_i)-e_f)}{L(t_i)}\theta(t_i)\right]<\varepsilon$$

$e_f$: length of focal exon of the skipped exon event ($e_2$)

$f$: set of focal transcript that includes the skipped exon event and was manipulated in simulation ($\{t_4\}$)

$ex$: set of other transcripts that exclude the focal exon ($\{t_2, t_3\}$)

$in$: set of other transcripts that include the focal exon ($\{t_1\}$)

$lfc$: log fold change for the manipulated transcript.

$\theta(t_i)$: base expression level of transcript $t_i$

$L(t_i)$: length of transcript $t_i$

Supplementary Figure S13. The condition that results in similar ratio of '*this*' and '*others*' after simulation of positive fold change.

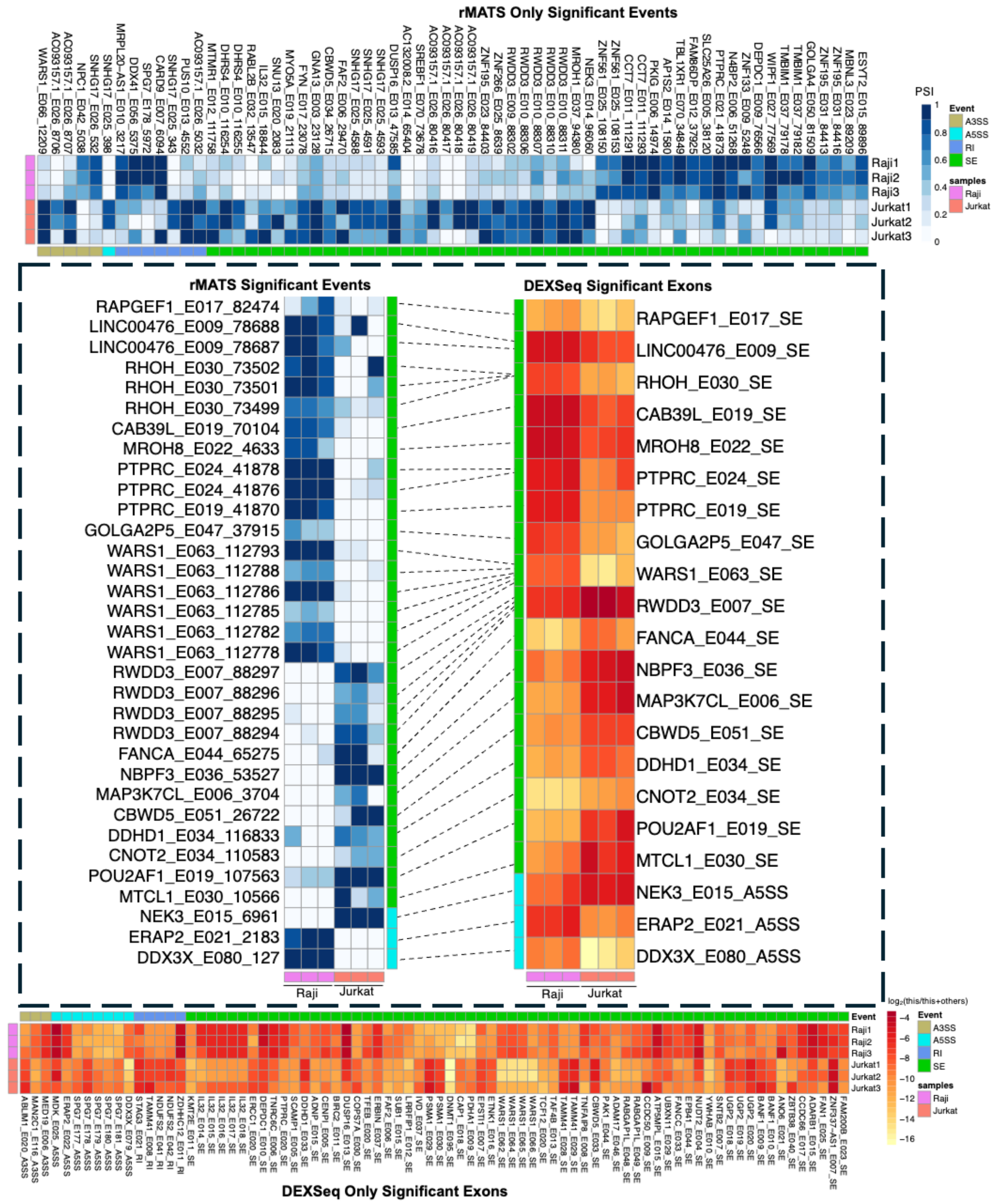


Supplementary Figure S14: Heatmaps of significant genes detected by rMATS and DEXSeq on Raji and Jurkat cell lines. The columns of the heatmap represent the genes grouped by the four AS events: khaki = alternative 3' splice site (A3SS), turquoise = alternative 5' splice site (A5SS), blue = retained intron (RI), green = skipped exon (SE). The rows represent the samples: violet = B naive cells, salmon = CD8$^{+}$ naive cells. The top heatmap display the inclusion levels (PSI) of the significant genes (FDR $\leq 0.05$) only detected by rMATS. Annotated on the columns are the gene symbols followed by the DEXSeq exon and then the unique rMATS ID. The bottom heatmap display the this/(this+others) counts $\log_2$ transformed of the significant genes (adjusted p-value $\leq 0.05$) only detected by DEXSeq. The gene symbols are followed by the DEXSeq exon and then the corresponding AS event. The middle heatmaps represent significant genes that are shared between rMATS (left) and DEXSeq (right) connected by dashed lines.
